# Revisiting PFA-mediated tissue fixation chemistry: *FixEL* enables trapping of small molecules in the brain to visualize their distribution dynamics

**DOI:** 10.1101/2021.12.21.473647

**Authors:** Hiroshi Nonaka, Takeharu Mino, Seiji Sakamoto, Jae Hoon Oh, Yu Watanabe, Mamoru Ishikawa, Akihiro Tsushima, Kazuma Amaike, Shigeki Kiyonaka, Tomonori Tamura, A. Radu Aricescu, Wataru Kakegawa, Eriko Miura, Michisuke Yuzaki, Itaru Hamachi

**Author notes:** These authors contributed equally to this work (Hiroshi Nonaka, Takeharu Mino). **Corresponding Author:** Prof. Itaru Hamachi, Department of Synthetic Chemistry & Biological Chemistry, Graduate School of Engineering, Kyoto University.

## Abstract

Various small molecules have been used as functional probes for tissue imaging in medical diagnosis and pharmaceutical drugs for disease treatment. The spatial distribution, target selectivity, and diffusion/extrusion kinetics of small molecules in structurally complicated specimens are critical for function. However, robust methods for precisely evaluating these parameters in the brain have been limited. Herein we report a new method termed “*Fix*ation-driven chemical crosslinking of *e*xogenous *l*igands (*FixEL*)” which traps and images exogenously administered molecules-of-interest (MOI) in complex tissues. This method relies on proteins-MOI interactions, and chemical crosslinking of amine-tethered MOI with paraformaldehyde used for perfusion fixation. *FixEL* is used to obtain images of the distribution of the small molecules and their dynamics, which addresses selective/nonselective binding to proteins, time-dependent localization changes, and diffusion/retention kinetics of MOI such as PET tracer derivatives or drug-like small molecules. Clear imaging of a nanobody distributed in the whole brain was also achieved with high spatial resolution using 2D/3D mode.

## Introduction

Tissues of living animals comprise various cells containing many of different molecules such as proteins, DNA/RNA, saccharides, and diverse small molecules. In the recent decade, the combination of hydrogel chemistry with tissues, termed hydrogel-tissue chemistry (HTC),^1–3^ has rapidly produced valuable methods capable of visualizing spatial distribution of these molecules in tissues with 3D manner, such as CLARITY,^4^ Sca*l*e,^5^ CUBIC,^6^ SeeDB,^7^ BABB,^8^ and 3DISCO^9^. These methods physically or chemically fix and entrap protein and RNA while retaining 2D/3D arrangement of tissues within a hydrogel matrix.^10–12^ The resultant hydrogel-tissue composites are visibly clarified; as a result, the fixed proteins/DNA can be visualized in the 3D mode of the whole tissue with advanced microscopy and conventional 2D microscopy. More recently, expansion microscopy using expandable synthetic hydrogels has provided nanometer-level spatial resolution for analysis of fixed biopolymers (proteins/DNA and RNA) in a tissue composite while maintaining the 3D distribution data.^13–17^ These methods are powerful; however, they do not apply well to the 3D analysis of small molecules because most of the small molecules cannot be effectively trapped in complex tissues using the present HTC technologies.

Various small molecules are used as functional probes for tissue imaging in medical diagnosis and as pharmaceutical drugs for disease treatment. These are exogenously administered to a live specimen and they act as a reporter and/or regulator to address the normal/abnormal state of a class of cells or tissues. Spatial distribution, target selectivity, and diffusion/extrusion kinetics of these molecules in structurally complicated specimens are critical to optimizing the function of small molecules. However, robust methods for precisely evaluating these parameters in tissues with high spatiotemporal resolution have been limited due to the structural complexity, which is in sharp contrast with the tools/methods developed for cultured cell analyses. A representative imaging method is positron emission tomography (PET), which can non-invasively visualize physiological and pathological changes *in vivo* through small molecule probes labeled with radioisotopes such as ^11^C and ^18^F. Although powerful, radiolabeled molecules are difficult to handle because of the radiation exposure and short lifetime.^18,19^ PET imaging requires a special instrument for measurement, and often suffers from low spatial resolution. Mass spectroscopy (MS) imaging is another useful method that detects the mass of small molecules in a tissue slice section without labeling.^20–22^ However, MS imaging usually requires complicated sample preparation protocols, lacks quantification, has low spatial resolution, and is difficult to obtain 3D information of the distribution of small molecules. As a result, the development of a simple and high-resolution 3D imaging method for small molecules in complicated tissues remains challenging.

Herein, we developed a new method termed “*Fix*ation-driven chemical crosslinking of *e*xogenous *l*igands (*FixEL*)”, which relies on interactions between proteins and small molecules and chemical crosslinking with paraformaldehyde (PFA), for imaging of exogenously administered molecules-of-interest (MOI) in complex tissues such as a mouse brain. *FixEL* has been developed by revisiting the traditional PFA-based tissue fixation and combining it with a novel chemical twist, which enables to obtain snapshot images of MOI in tissues. Because *FixEL* is also highly compatible with most tissue clearing technologies, as well as conventional immunostaining of sliced tissues, we are able to address the selective/nonselective interactions to proteins, time-dependent localization changes, and diffusion/retention kinetics of exogenously-added MOIs such as PET tracer derivatives and drug-like small molecules. Additionally, the spatial distribution of a nanobody in the 2D/3D mode of the whole brain was visualized with a high resolution of the synaptic puncta using *FixEL*.

## Results

### Strategy of *FixEL* in live tissues

Perfusion fixation using PFA (formaldehyde) aqueous solution was combined with chemical crosslinking of MOIs in the *FixEL* method. The formaldehyde-based tissue fixation conducted by soaking or perfusion originated in the 1890s by Ferdinand Blum *et al*.^23,24^ Nowadays, perfusion fixation is widely used in biology and pathology to fix tissues/cells of biological samples while maintaining the 2D/3D space.^25,26^ PFA fixation employs an aqueous formaldehyde solution consisting of polymeric formaldehyde with various degree of polymerization (Figure 1a). Aldehyde moieties of both terminals of PFA can readily react with the nucleophilic side chains (such as lysine) of many proteins in biological samples. Because most natural proteins have multiple nucleophilic amino acids and PFA is a bifunctional reactive polymer, PFA fixation produces a hydrogel-like 3D construct where PFA-modified proteins are efficiently fixed as crosslinking points (Figure 1a). However, most small molecules are not effectively trapped in this hydrogel matrix because they do not have such nucleophilic reactive groups and the mesh size of the resultant hydrogel is too large to entrap such small molecules. We thus decided to introduce a primary amine to a small molecule-of-interest (sMOI), to serve as a nucleophile that is reactive with PFA. Additionally, we hypothesized that attractive interactions of sMOI with proteins may assist rapid/effective crosslinking with PFA, which allows for immobilizing sMOI covalently near interacting proteins during PFA-mediated fixation, as shown in Figure 1a.

**Figure 1.**
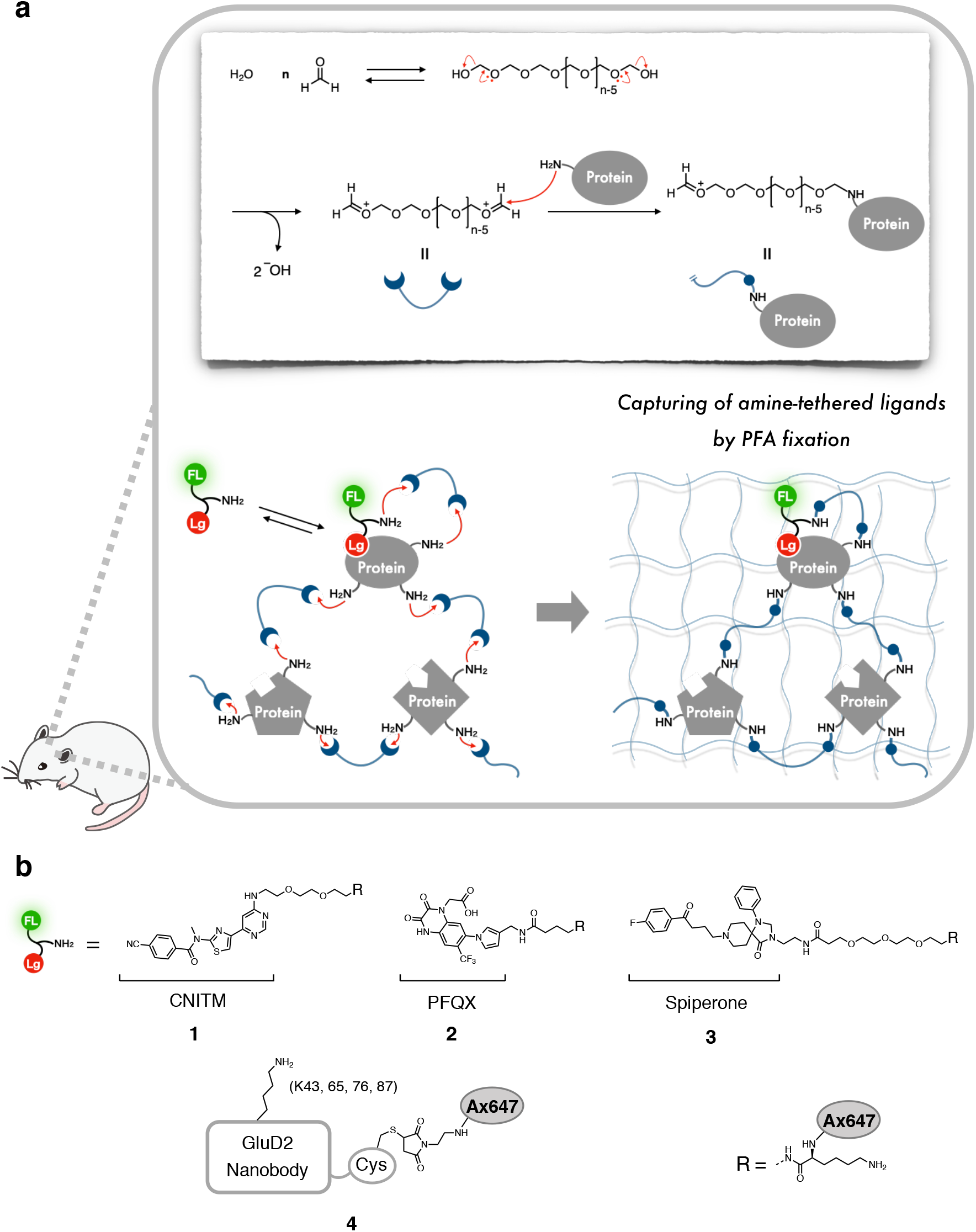
Fixation-driven chemical crosslinking of exogenous ligands (*FixEL*) in the mouse brain. **a**, Schematic illustration of the *FixEL* strategy. FL and Lg refer to fluorophore and ligand, respectively. **b**, *FixEL* probes used in this study. **1** has a CNITM moiety, which is a ligand for mGlu1, and an amino group. **2** has a PFQX moiety, which is a ligand for AMPAR, and an amino group. **3** has a Spiperone moiety, which is a ligand for DRD2, and an amino group. **4** is an Ax647-conjugated GluD2 nanobody.

A few ligand molecules that are known to selectively interact with a particular protein were tested as the sMOI and modified with a reactive amino group and fluorescent reporter for imaging analysis in tissues (Figure 1b). When a modified sMOI is exogenously administered to a live mouse, it reversibly interacts with a protein to form a transient complex. Upon infusion of PFA into the specimen, the amino group of sMOI are crosslinked with a nucleophilic amino acid of interacting proteins through hemiaminal bonds of PFA, resulting in covalent entrapment of the modified sMOI in the PFA-hydrogel matrix. Perfusion fixation using PFA achieved rapid and uniform immobilization of tissues, which allowed for the capture of the living state of a specimen with minimal risk of death-induced autolysis. Subsequently, the resultant PFA-fixed tissues containing sMOI can be subjected to several analytical workflows, such as fluorescent microscopy imaging and immunostaining for studying spatial localization of sMOI in sliced 2D tissues and for evaluating the 3D distribution of sMOI in a mouse brain using transparent samples prepared by tissue clearing protocols. Additionally, varying the initiation time of the PFA perfusion can be used to fix distinct states of a specimen, whose tissue imaging provided valuable snapshot data, such as the time-dependent distribution of sMOI, that can be correlated with its diffusion and retention rate in the whole brain.

### Proof-of-principle experiments of *FixEL* in cultured cells

As a proof-of-principle study, the process was initiated with 4-cyano-*N*-[4-(6-(isopropylamino)-pyrimidin-4-yl)-1,3-thiazol-2-yl]-*N*-methylbenzamide (CNITM). CNITM is a derivative of a PET probe for metabotropic glutamate receptor type 1 (mGlu1),^27^ a type-C GPCR expressed in the cerebellum and thalamus regions, whose expression level may be related to Parkinson’s disease.^28,29^ CNITM bearing the affinity with mGlu1 (*K*_i_ = 27 nM), was modified with a primary amine and a fluorescent AlexaFlour647 (Ax647) dye to produce the CNITM Probe **1** (Figure 1b). The accessible site for such modification was judged from the co-crystal structure of mGlu1 and FITM, a ligand similar to CNITM.^30^ Three control molecules shown in Figure 2a were also prepared.

**Figure 2.**
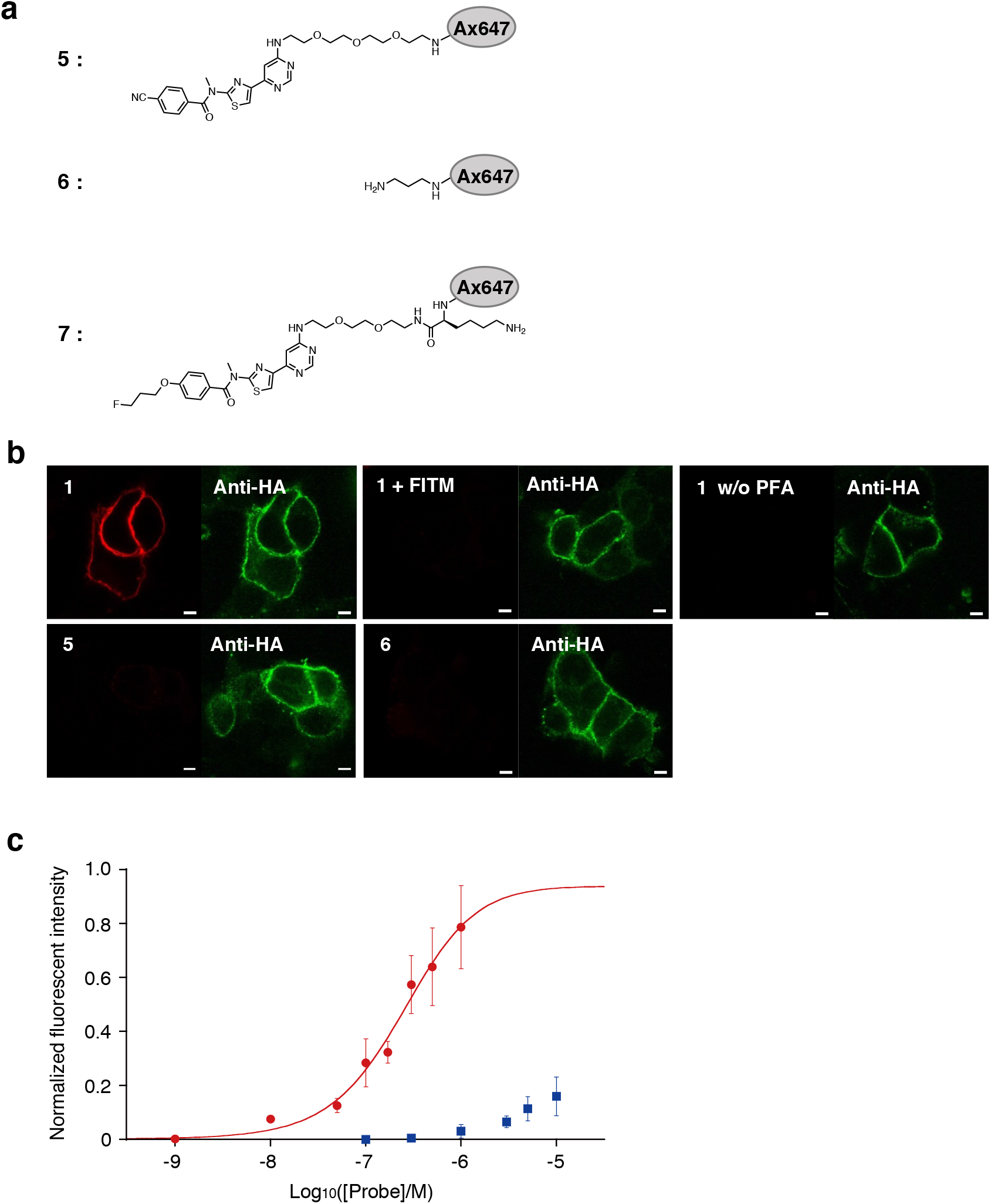
*FixEL* in cultured cells. **a**, Chemical structures of control molecules **5**-**7**. **b**, Confocal imaging of HA-tag fused mGlu1 expressed on HEK293T after *FixEL* of probe **1**, **1** with 20 eq. of FITM, **1** without PFA treating, **5**, or **6** (Ax647, red). HA-tag fused mGlu1 was stained with DyLight550 anti-HA tag (green). Fluorescence imaging of the cells was performed using a CLSM equipped with a 63× objective and a GaAsP detector (561 nm excitation for DyLight550 and 633 nm excitation for Ax647). Scale bar 5 μm. **c**, Fluorescence change of the plasma membrane on mGlu1-expressed HEK293T cells depending on the concentration of probe **1** (red circle) or probe **7** (blue square). n = 8 cells, Data are presented as mean ± s.e.m.

*FixEL* experiments were first conducted using HEK 293T cells expressing mGlu1 (Figure 2b). After incubation of mGlu1-expressing cells with probe **1** for 5 min, the cells were treated with PFA (30 min), followed by washing with DMEM (37°C, overnight). The fluorescence of Ax647 was observed from the plasma membrane of mGlu1-expressing cells using confocal laser scanning microscopy (CLSM). Such fluorescence was not observed with probe **5** that lacks the amine moiety, clearly indicating the reactive amine is essential for this crosslinking. Under the conditions where FITM bearing the stronger affinity than CNITM was co-incubated with **1**, the fluorescence was nearly diminished. Negligible fluorescence was detected in the case of probe **6** with an amine and Ax647, and no appropriate ligand. Therefore, the CNITM-mGlu1 interaction is also essential for the crosslinked fixation of **1**. No addition of PFA did not create fluorescence at the plasma membrane of HEK 293T cells (Figure 2b). Therefore, the probe **1** fixation was carried out by the amine-originated crosslinking of PFA with essential assistance of the protein–ligand interaction. In the titration experiment with varied concentration of probe **1**, the fluorescence intensity from the plasma membrane showed a typical saturation behavior with 243±77 nM of *K*_d_, whose value was nearly identical with CNITM, confirming the binding of probe **1** with mGlu1 is a key control factor in this crosslinked fixation (Figure 2c). In the case of **7** with a lower affinity ligand (*K*_i_ > 5 μM),^31^ the fluorescence intensity from the plasma membrane did not show saturation behavior up to 10 μM, indicating that *FixEL* can reflect the affinity of MOI to a binding protein.

### *FixEL* of the small molecule probe for mGlu1 in a mouse cerebellum

*FixEL* was used in a mouse cerebellum whose molecular layer highly expressed endogenous mGlu1, to examine the spatial distribution of **1**. Probe **1** in an aqueous solution was injected directly into the cerebellum of 5-week-old live C57BL/6N mice under anesthesia; thereafter, we confirmed the mice were alive and freely moved. Perfusion fixation by PFA was performed 14 h after the probe injection, and the fixed cerebellum was cut into 50-μm thick sections by cryostat (Figure 3a, Supplementary Figure 1). CLSM observation of the sections showed that Ax647-derived fluorescence was observed selectively in the molecular layer region, which was in good agreement with the expression area of mGlu1 (Figure 3b).^32^ Such clear and selective fluorescence was not observed in the cases of probes **5** or **6** that lack either the amine group or ligand part, respectively. Additionally, in the experiment using mGlu1 KO mice, the fluorescence signal was strongly diminished, which suggests that probe **1** was fixed proximal to mGlu1, and both the amine tethering and ligand–protein interaction are essential for the effective entrapment of **1**.

**Figure 3.**
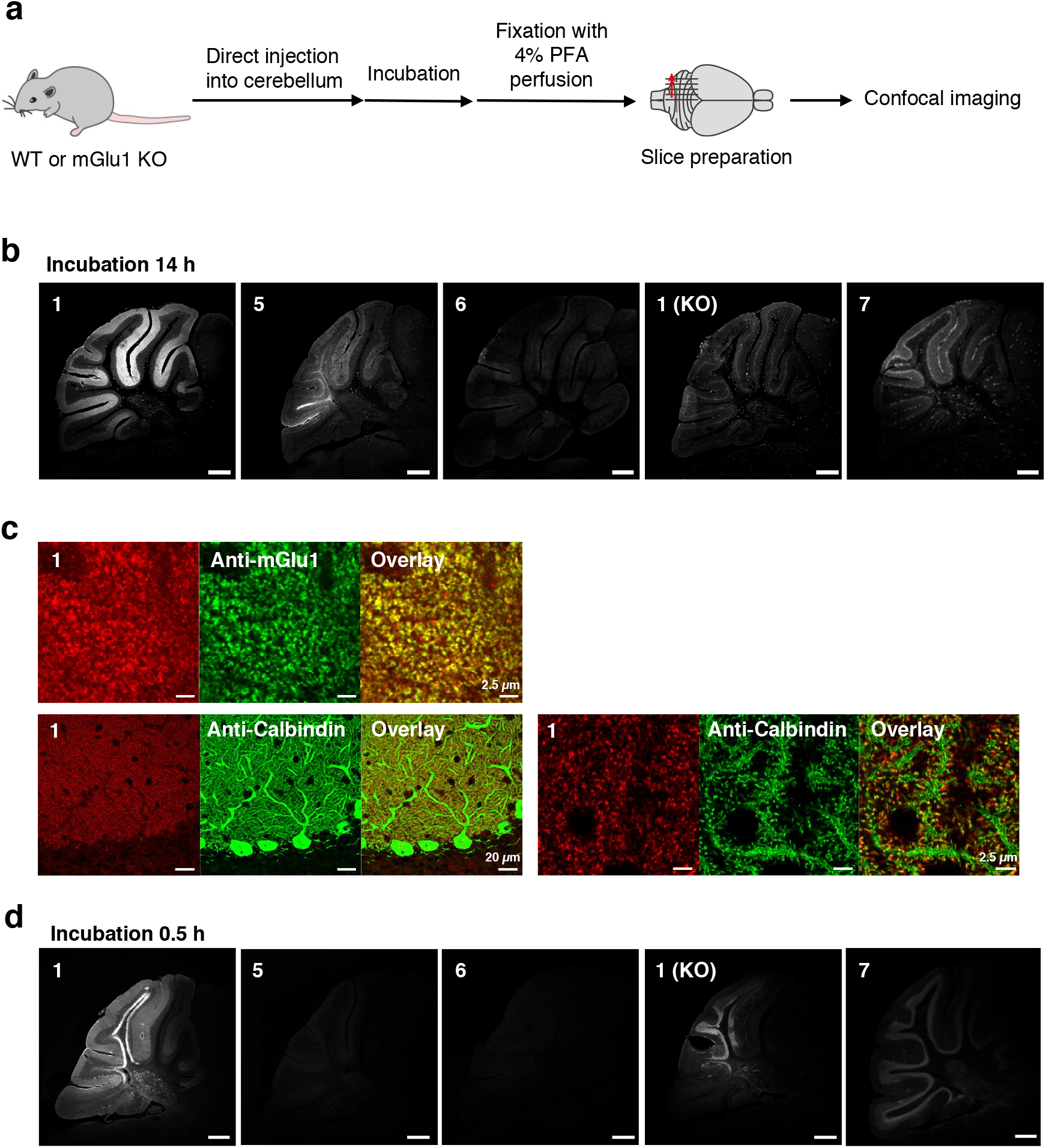
*FixEL* of a small molecule ligand for mGlu1 in the mouse cerebellum. **a**, Schematic illustration of experimental procedures. PBS(−) containing probe **1** (4.5 μL of 10 μM probe which is assumed to be ca. 0.7 μM (C_cere_) on the basis of the cerebellum volume^48^) was injected into mouse cerebellum. After 0.5 h or 14 h of incubation, the mouse was transcardially perfused with 4% PFA. The cerebellum was isolated and sectioned by cryostat (50-μm thick). **b**, Fluorescence imaging of cerebellum slices with *FixEL* of probe **1** after 14h incubation. Imaging was performed using a CLSM equipped with a 10× objective and GaAsP detector (633 nm excitation for Ax647). From left to right images, probes **1**, **5**, **6**, and **1** in mGlu1 KO mouse, and **7** were used. Scale bar: 500 μm. **c,** Co-immunostaining of the cerebellum slice with *FixEL* of probe **1** after 12h incubation. Probe **1** is shown in red, and anti-mGlu1 and anti-calbindin are shown in green. The slices after *FixEL* of probe **1** were permeabilized and immunostained with primary antibody anti-mGlu1 or anti-calbindin and secondary antibody (Alexa Fluor 488(Ax488)) in 0.1% Triton X-100/PBS(−). Fluorescence images in top left and bottom right were acquired by using a CLSM equipped with a 100× objective, GaAsP detector (488 nm excitation for Ax488 and 633 nm excitation for Ax647), and Lightning deconvolution. Fluorescence image in top right was acquired by using a CLSM equipped with a 63× objective and GaAsP detector (488 nm excitation for Ax488 and 633 nm excitation for Ax647). **d**, Fluorescence imaging of cerebellum slices with *FixEL* of probe **1** after 0.5h incubation. Imaging was performed using a CLSM equipped with a 5× objective. Scale bar: 500 μm.

High-resolution CLSM imaging confirmed that the fluorescence in the molecular layer comprised an assembly of many bright puncta with μm size, which is consistent with the perisynaptic localization of mGlu1 (Figure 3c). The dendritic spines derived from Purkinje cells were stained with anti-Calbindin, a Purkinje cell marker. Immunostaining of mGlu1 showed that many of the bright spots of Ax647 merged well with the fluorescence from anti-mGlu1. All of these results strongly demonstrate that the location of the CNITM probe **1** was visualized by *FixEL*, and **1** was retained proximal to mGlu1 in the molecular layer of the cerebellum through CNITM-mGlu1 selective interactions for at least 14 h in the live mouse.

The short incubation time (0.5 h) after the probe **1** injection, on the other hand, produced a nonselective image with strong fluorescence near the injection site, in which the fluorescent areas were broadly spread in the granular layer and white matter, other than the molecular layer (Figure 3d). This nonselective fluorescence may be explained by that an excess amount of **1** remained in the injection area in an early time range after the injection and **1** can be fixed by PFA perfusion through nonspecific interactions with proteins other than mGlu1. Given that such images were not obtained for the control probes **5** and **6**, the nonspecific interactions reflect the distinct physicochemical properties of the ligand including the hydrophobicity and presence/absence of the tethered amine. Such nonspecific fluorescence was reduced after 14 h incubation, and selective fluorescence of the molecular layer clearly appeared. In the case of probe **7**, a short incubation time (0.5h) produced weak fluorescence in the granular layer and Purkinje cell bodies and after the prolonged incubation (14 h), a non-specific signal similar to KO sample was observed, indicating the poor targetability and lower retention of this weaker affinity ligand.

The diffusion behavior of probe **1** was visualized (Supplementary Figure 1) using the distance-dependent imaging data. The perfusion fixation was performed 12 h after the injection, followed by slices 0.6, 1.2, and 1.8 mm from the right side of the injection point, the cerebellar vermis, 0 mm. Selective fluorescence of the molecular layer was observed in all slices, while the signal intensity gradually decreased depending on the distance from the injection point, which implies that probe **1** penetrated, diffused and retained along the direction away from the injection site and a concentration of **1** reduced around the edges of the cerebellum.

### *FixEL* to visualize sMOI of mGlu1 in the whole brain

The direct injection of probe **1** from the lateral ventricle (LV) allowed for analyzing its spatial distribution in the whole brain (Figure 4a and 4b). Various molecules, such as peptides, proteins, and DNA/RNA, injected into LV are diluted by the cerebrospinal fluid and diffuse with its flow. After injection of probe **1** into the LV, perfusion fixation with 4% PFA was performed with incubation times of 0.5, 1, 3, 6, 12, and 24 h. Sections 600 μm from the brain midline were prepared and observed using CLSM (Figure 4a). mGlu1 is expressed in the thalamus and olfactory bulb, as well as the cerebellum of the whole-brain. For the slice fixed 0.5 h after injection, strong fluorescence was observed near the LV region and at the outer edge of the cerebellum where the cerebrospinal fluid is in contact. The slice after 1 h incubation showed widely distributed fluorescence from the LV region to the thalamus and midbrain areas, suggesting that the probe penetrated into the brain parenchyma and was trapped there through crosslinking with nonspecific interactions caused by the high concentration of **1**, whereas the fluorescence derived from the selective interaction to mGlu1 became clearer in the cerebellar molecular layer. In the slice after 3h of incubation, the overall fluorescence intensity was less than shorter incubations, probably because of the washing-out effect with cerebrospinal fluid. Additionally, the selective staining of the molecular layer in the cerebellar region was more clearly visualized while the fluorescence due to nonspecific interactions still remained in the mid-brain areas. The fluorescence reflecting the mGlu1 expression areas in the molecular layer of the cerebellum and thalamus gradually faded 6–24 h after injection, whereas the fluorescence in others areas nearly disappeared at 6 h. This suggests the probe fixed due to nonspecific interactions was more rapidly excreted with the flow of cerebrospinal fluid. The snapshot images with high-resolution allowed us to chase the diffusion of probe **1** in more closely (Figure 4b). At the edge of the cerebellar molecular layer that is the contact surface with the cerebrospinal fluid, the fluorescence strongly appeared 0.5 h after injection and intensified over 3 h. The fluorescence at the interface between the molecular layer and granular layer was weak relative to that of the contact area, resulting in a fluorescence gradient from the contact area to the inner interface. The gradient gradually became gentle and flat over 12 h. This implied that probe **1** permeated from the contact area, diffused to the molecular layer, and distributed homogenously in the molecular layer; however, probe **1** was not trapped in the granular layer; thereafter, the probe was slowly discharged. These results suggest that *FixEL* is a unique technique capable of visualizing the diffusion dynamics of sMOI, as well as the spatial distribution of sMOI in the whole brain with high resolution.

**Figure 4.**
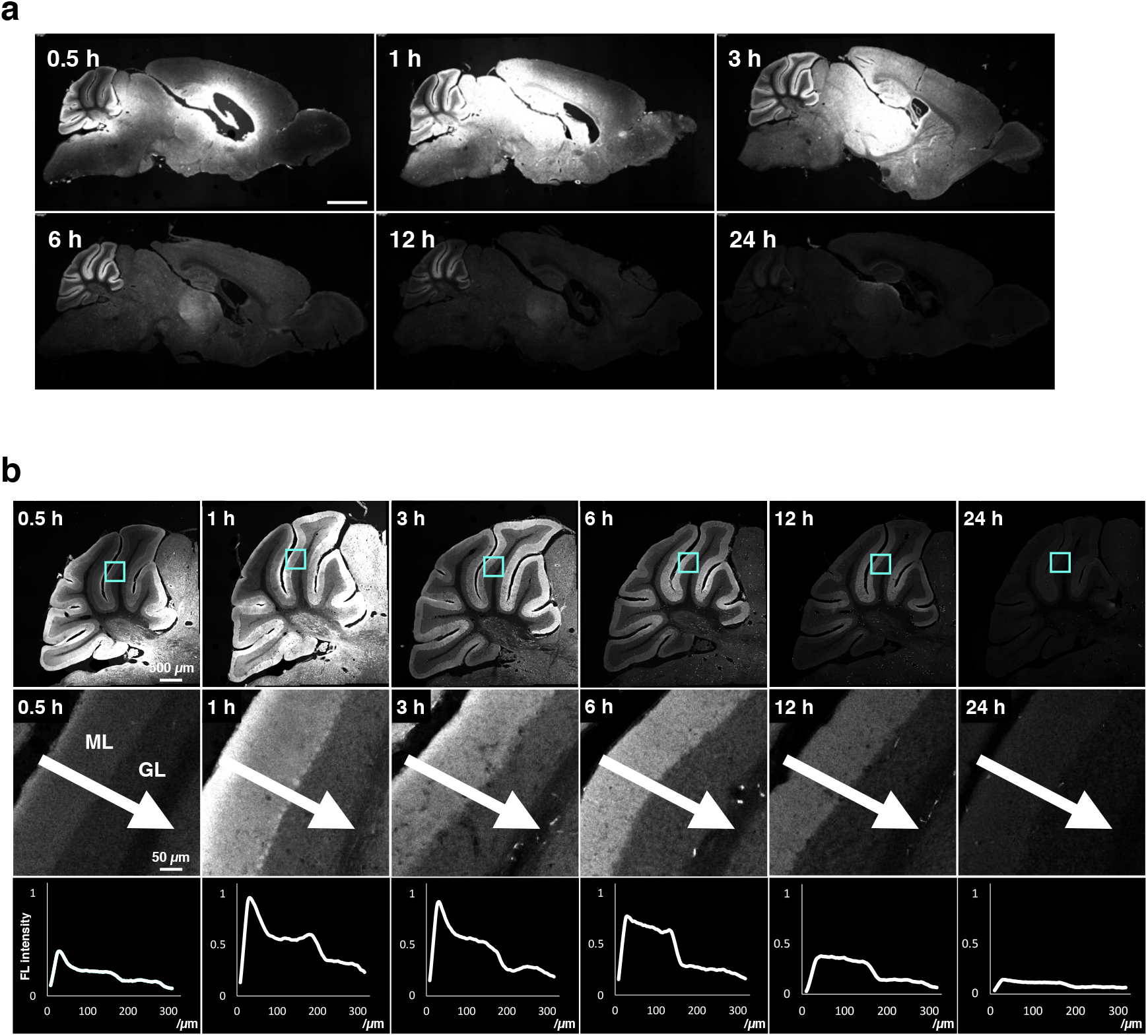
Visualization of the distribution dynamics of probe 1 after *FixEL* at different incubation times. **a**, Fluorescence imaging of whole-brain sagittal slices after *FixEL* with probe **1**. PBS(−) containing probe **1** (4.5 μL of 100 μM probe which is assumed to be ca. 1 μM (C_brain_) on the basis of the brain volume^48^) was injected into the mouse lateral ventricle. After 0.5, 1, 3, 6, 12, or 24 h of incubation, mouse was transcardially perfused with 4% PFA. Fluorescence imaging of slices (50-μm thick) was performed using a CLSM equipped with a 5× objective, and GaAsP detector (633 nm excitation for Ax647). Scale bar: 2 mm. **b**, Line plot analysis of distribution dynamics of probe **1** in the cerebellar lobule V after injection into lateral ventricle. Fluorescence imaging of the cerebellum region was performed using a CLSM equipped with a 10× objective. The light blue square region was magnified. The line plot analysis was performed along a white arrow.

### *FixEL*-based imaging of various MOI probes in the whole brain

*FixEL* was applied to other small molecular ligands, such as 6-pyrrolyl-7-trifluoromethyl-quinoxaline-2,3-dione (PFQX) and spiperone to demonstrate the robustness of the method. PFQX is a selective antagonist of the AMPA receptor, an ionotropic glutamate receptor.^33–35^ According to the design guideline of probe **1** (for mGlu1), a PFQX-tethered probe **2** was chemically synthesized (Figure 1b). After direct injection of probe **2** into LV and perfusion fixation, the slice samples were prepared. Fluorescence signals were mainly observed in the hippocampus, cerebellum, and cortex where AMPA receptors are naturally expressed (Figure 5a). We confirmed that the fluorescence signal of probe **2** merged with the immunostaining signal of anti-GluA2 (Supplementary Figure 2a). Spiperone, which has been used as psychiatric drug, is an antagonist of the D2 dopamine receptor and is its PET tracer.^36–38^ We similarly synthesized a spiperone-tethered probe **3** (Figure 1b). The fluorescence signals of the slice prepared by the *FixEL* protocol were clearly observed in the striatum where endogenous D2 receptors are expressed (Supplementary Figure 2b). We also detected fluorescence in the middle of the cerebral cortex. Because spiperone is also known to bind to the 5HT2A serotonin receptor expressed in the cerebral cortex, this fluorescence might be derived from the spiporone-5HT2A interaction (*K*_i_ for DRD2 = 0.06 nM, *K*_i_ for HT2A ≈ 1–2 nM).^39–41^ The fluorescence signal of probe **3** merging with the immunostaining signal was confirmed using anti-DRD2 in striatum region and 5HT2A in cerebral cortex region (Figure 5b), demonstrating *FixEL* provided the ability to image the distribution of sMOI relying on the interaction with a target protein, and the off-target protein in the brain.

**Figure 5.**
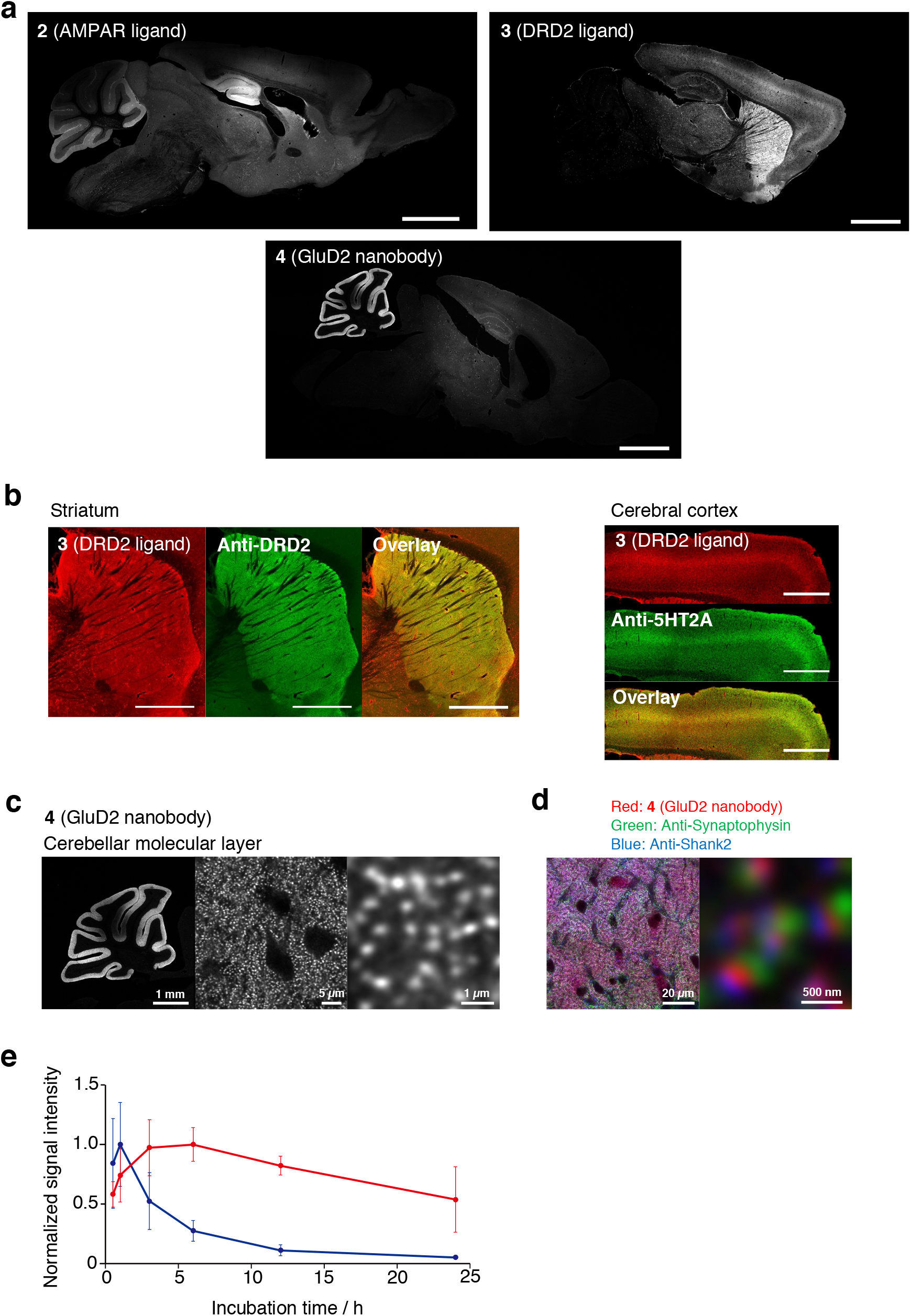
Distribution and localization analysis of *FixEL* probes 2–4 in the whole brain. **a**, Fluorescence imaging of whole-brain sagittal slices after *FixEL*. PBS(−) containing a probe (4.5 μL) was injected into mouse lateral ventricle. Conditions: 40 μM **2** (C_brain_ is ca. 0.4 μM), incubation time 3 h. 25 μM **3** (C_brain_ is ca. 0.25 μM), incubation time 16 h. 40 μM **4** (C_brain_ is ca. 0.4 μM), incubation time 24 h. Fluorescence imaging of slices (50-μm thick) was performed using a CLSM equipped with a 10× or 5× objective and GaAsP detector (633 nm excitation for Ax647). Scale bar: 2 mm. **b**, Co-immunostaining of the slices after *FixEL* with probe **3**. The slices were fixed, permeabilized and immunostained using anti-DRD2 or anti-5HT2A (green, Ax488-conjugated secondary antibody). Scale bar: 1 mm. **c**, Higher magnification image of the sample in Figure 5a stained with probe **4**. Fluorescence imaging was performed using a CLSM equipped with a 10× objective (left panel) and a 100× objective with Lightning deconvolution (middle and right panel). **d**, Co-immunostaining of the sagittal slices with *FixEL* probe **4**. The slices after *FixEL* were permeabilized and immunostained using anti-synaptophysin (green, Ax488-conjugated secondary antibody) and anti-Shank2 (blue, Ax405-conjugated secondary antibody). Fluorescence imaging was performed using a CLSM equipped with a 100× objective, a GaAsP detector (405 nm excitation for Ax405, 488 nm excitation for Ax488, and 633 nm excitation for Ax647), and Lightning deconvolution. **e**, Comparison of the time-dependent decrease in fluorescence of cerebellar regions between slice samples with *FixEL* using probe **1** (blue circle) and probe **4** (red circle). Average of four ROIs. Data are presented as mean ± s.e.m.

As MOI other than small molecules, GluD2-Nanobody (15 kDa), a down-sized antibody that binds to a glutamate receptor GluD2 (Figure 1b), was evaluated.^42,43^ GluD2 is endogenously expressed at the postsynapse of the cerebellum molecular layer and it functions as a bidirectional synaptic organizer through a triadic complex formation with neurexin and Cbln1.^44^ Because the GluD2 nanobody contains a few reactive Lys amino groups that are useful for fixation-driven crosslinking, we simply needed to modify the nanobody with Ax647 (Ax647-GluD2 Nb shown in Figure 1b). Ax647-GluD2 Nb (probe **4**) was injected into LV of 5-week-old C57BL/6N mice followed by perfusion-fixation and the slice samples were prepared, according to the *FixEL* protocol. Strong fluorescence signal was observed predominantly in the cerebellar molecular layer of the slices (Figure 5a) that merged well with anti-GluD2 (Supplementary Figure 2c). The signal comprised many small bright puncta smaller than 1 μm (Figure 5d). The immunostaining experiments of this slice using anti-Synaptophysin, a presynaptic marker, and anti-Shank2, a postsynaptic marker, showed that the puncta of probe **4** merged well with those of anti-Shank2, and neighbored the signals of anti-Synaptophysin (Figure 5d). These results demonstrated that the GluD2 Nb interacts with the GluD2 in a narrow 20–30 nm synaptic cleft of the molecular layer of the cerebellum. Overall, each tested MOI showed the distinct spatial distribution dependent on the expression areas of proteins targeted by the corresponding ligands, which indicates the chemical decoration of the ligand with an amine and fluorescent Ax647 may have minimal impact on the corresponding ligand properties.

Imaging was conducted to reveal the diffusion dynamics of the GluD2 nanobody in the whole brain of a mouse. As shown in Supplementary Figure 3, clear fluorescence was observed in the cerebellar molecular layer and areas near the injection site, which were attributed to GluD2 selective binding and nonspecific interactions, respectively, in the early stages. The images from an incubation longer than 12 h indicated that the nanobody bound via nonselective interactions were excreted more consistently, whereas nanobody selectively bound to GluD2 remained tightly bound. It is also interesting to compare the retention kinetics in the cerebellum between the CNITM probe **1** and GluD2 nanobody **4** (Figure 5e). The time profile of the fluorescence intensity of Ax647-GluD2 nanobody showed a biphasic behavior including the fluorescence increase in the initial 0.5–6 h and subsequent decrease, indicating that nanobody penetrated/condensed up to 3–6 h and then was slowly excreted with a half-life of longer than 24 h. On the other hand, CNITM probe **1** showed a rather rapid increase (within 1 h) and decrease with a half-life of approximately 6 h. This difference in CNITM and GluD2 nanobody retention time in the cerebellum molecular layer may be attributed to the dissociation rate of the corresponding ligand–protein interaction (i.e., CNITM-mGlu1 vs nanobody-GluD2). We separately confirmed the much more rapid dissociation CNITM from mGlu1, compared to that of Nanobody from GluD2 using the live cell experiments (Supplementary Figure 4). The results indicate that the *FixEL* method has the potential to directly evaluate parameters related to the diffusion dynamics of MOIs in biological tissues.

### Transparent tissues prepared by combining *FixEL* and 3DISCO

The fixed tissue prepared by *FixEL* with probe **1** was compatible with the tissue clearing protocol of 3DISCO.^9^ The cerebellum tissue was made transparent by 3DISCO and the resultant tissue sample was subjected to 3D CLSM imaging, showing that **1** was selectively and three-dimensionally distributed in multiple molecular layers throughout the whole cerebellum (Supplementary Figure 5). A whole brain sample prepared by the LV injection of **1** was also clarified and 3D CLSM imaging was performed (Figure 6a–d). Clear fluorescence was observed in the thalamus and cerebellar molecular layer where mGlu1 was localized (Figure 6b, 6c). When the cerebellar molecular layer of this transparent sample was observed with a higher magnification lens of CLSM, the layer was observed to be densely packed with bright puncta smaller than 1 μm, likely derived from dendritic spines (Figure 6d). To demonstrate the versatility of 3D distribution analysis by combining *FixEL* and 3DISCO, we also performed 3D imaging of a whole brain treated with probe **4**. As shown in Figure 6e and 6f, the distribution of GluD2 nanobody **4** was clearly observed in the brain-wide level, revealing **4** was localized with remarkably high selectivity in the molecular layer (comprising many bright puncta) of the cerebellum where GluD2 was expressed. The 3D-distributions of PFQX probe **2** and spiperone probe **3** were also visualized, which agrees with the expression areas of the corresponding natural receptors (Supplementary Figure 6). Therefore, the combination of *FixEL* and tissue clearing technology such as 3DISCO, enabled the evaluation of the 3D distribution of MOIs in the brain from whole brain (mm) scale to dendritic spine (μm) scale.

**Figure 6.**
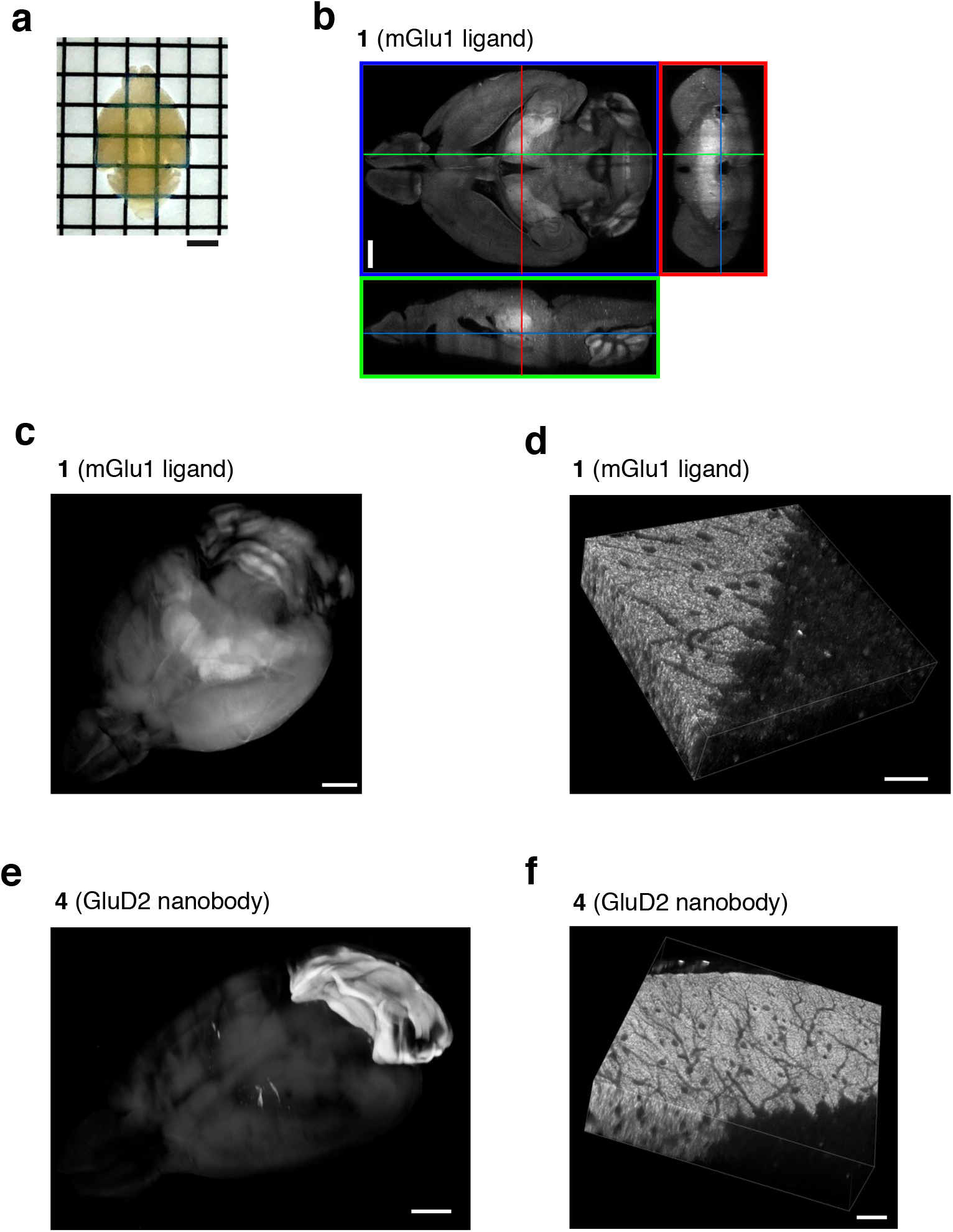
3D fluorescence imaging of *FixEL* samples with 3DISCO. **a**, Appearance of a brain sample after *FixEL* with probe **1** and 3DISCO. Scale bar 2.5 mm. **b**, Z-stacking fluorescence imaging of the Figure 6a sample. PBS(−) containing probe **1** (4.5 μL, 100 μM: C_brain_ is ca. 1 μM) was injected into mouse lateral ventricle. After 6 h of incubation, the mouse was transcardially perfused with 4% PFA. After 3DISCO treating, z-stacking fluorescence imaging of the whole brain was performed using a CLSM equipped with a 5× objective and GaAsP detector (633 nm excitation for Ax647). Scale bar: 1 mm. **c**, 3D rendering of the whole brain in Figure 6b. Scale bar: 1 mm. **d**, Z-stacking fluorescence imaging of cerebellum region in Figure 6c with a 40× objective. Scale bar: 20 μm. **e**, 3D rendering of the whole brain after *FixEL* with GluD2 nanobody **4** and 3DISCO. PBS(−) containing probe **4** (10 μM, 4.5 μL ×2: C_brain_ is ca. 0.2 μM) was injected into the mouse bilateral ventricle respectively. After *FixEL* (20 h incubation) and 3DISCO treating, z-stacking fluorescence imaging of the whole brain was performed using a CLSM equipped with a 5× objective and GaAsP detector (633 nm excitation for Ax647). Scale bar: 1 mm. **f,** Z-stacking fluorescence imaging of the cerebellum region of Figure 6e with a 40× objective. Scale bar: 20 μm.

## Discussion

By revisiting traditional PFA-mediated tissue fixation chemistry and coupling with rational molecular design, we were able to develop a new method to capture and visualize exogenously-administered small molecules in the brain. *FixEL* is rapid and simple method that does not require special instruments, while *FixEL* requires chemical modification of appropriate probes. Unlike PET imaging, *FixEL* cannot be used for real-time imaging of a live brain. Instead, imaging of a particular time point, i.e., a snapshot, was achieved using perfusion fixation of a live mouse with much higher spatial resolution. This method is highly compatible with existing analytical methods including CLSM, immunostaining and tissue clearing technology. *FixEL* thus can provide various information of MOI probes in a live brain, including target/off-target proteins binding in various areas, time-dependent localization changes, and diffusion/retention kinetics with high spatial resolution of the whole brain. We demonstrated the visualization of diffusion dynamics of PET tracer derivatives and drug-like small molecules, which is valuable for the development/optimization of receptor-targeting drugs and PET tracers. We also succeeded in imaging a nanobody location with the 2D/3D mode of μm-spatial resolution. To the best of our knowledge, this is the first report to reveal the selective distribution dynamics of a nanobody in a whole mouse brain, which may provide crucial and fundamental information for future applications of nanobody-based drugs and imaging probes. *FixEL* should be complementary to conventional PET imaging and IR imaging for small molecules in complex tissues and brain.

## Methods

### Synthesis

All synthesis procedures and characterizations are described in the Supplementary Information.

### Subcloning and preparation of a dye-conjugated GluD2 nanobody

The protocols of subcloning, preparation, and characterization of an Ax647-conjugated GluD2 nanobody are described in the Supplementary Information.

### Fluorescence imaging

Fluorescence imaging was performed using a CLSM (Leica microsystems, Germany, TCS SP-8) equipped with 5× objective (NA = 0.15 dry objective), 10× objective (NA = 0.40 dry objective), 40× objective (NA = 1.30 oil objective), 63× objective (NA = 1.40 oil objective), 100× objective (NA = 1.40 oil objective), and GaAsP detector. The excitation laser was derived from a white laser and was set to an appropriate wavelength depending on the dye. Lightning deconvolution process (LAS X 3.5.5, Leica microsystems, Germany) was used in Figure 3c, 5c, and 5d. The procedures and conditions are described in the following figure captions and the Supplementary Information.

### Injection of reagents into the mouse cerebellum

Experiments were conducted according to the literature using 5 weeks old mice (male, C57BL/6N strain; body weight 18–22 g).^45^ mGlu1 KO mice were purchased from Laboratory Animal Resource Center (Tsukuba University, Japan).^46^ Under the deep anesthesia, probe **1** (4.5 μL) was directly injected into the vermis of cerebellar lobules V-VIII (0.5 mm depth from the surface) using a microinjector (Nanoliter 2010, world precision instruments) (600 nL/min).

### Injection of reagents into mouse lateral ventricle

Experiments were conducted according to the literature using 5 weeks old mice (male, C57BL/6N strain; body weight 18–23 g).^47^ Under the deep anesthesia, the probe or nanobody solution (4.5 μL) was directly injected into the lateral ventricle using a microinjector (Nanoliter 2010, world precision instruments) (600 nL/min).

### Animal experiments

C57BL6/N mice were purchased from Japan SLC, Inc (Shizuoka, Japan). The animals were housed in a controlled environment (23 °C, 12 h light/dark cycle) and had free access to food and water, according to the regulations of the Guidance for Proper Conduct of Animal Experiments by the Ministry of Education, Culture, Sports, Science, and Technology of Japan.

All experimental procedures were performed in accordance with the National Institute of Health Guide for the Care and Use of Laboratory Animals, and were approved by the Institutional Animal Use Committees of Kyoto University.

## Author contributions

I.H. conceived and designed the project. T.M., J.H.O., M.I., A.T., and K.A. performed the probe syntheses and preparation. T.M., S.S., H.N., S.K., and K.A. performed the imaging experiments related to probe **1**. H.N., S.K., and K.A. performed the experiments related to PFQX probe **2**. H.N. and A.T. performed the experiments related to Spiperone probe **3**. J.H.O., H.N., Y.W., and T.T. performed the experiments related to GluD2 nanobody **4** with the help of A.R.A.. H.N., T.M., S.S., J.H.O., A.T. and K.A. performed animal experiments with the help of S.K., E.M., W.K., and M.Y.. H.N., T.M., S.S., and I.H. wrote the manuscript. All authors discussed and commented on the manuscript.

## Acknowledgements

The authors thank Dr. Lei Wang and Dr. Muneo Tsujikawa for organic synthesis and technical supports of biological experiments. The authors also thank Ashleigh Cooper, PhD, from Edanz (https://jp.edanz.com/ac) for editing a draft of this manuscript. This work was funded by the Japan Science and Technology Agency (JST) ERATO Grant No. JPMJER1802 and by a Grant-in-Aid for Scientific Research on Innovative Areas “Chemistry for Multimolecular Crowding Biosystems” (JSPS KAKENHI Grant No. 17H06348) to I. H.

## Data availability

The authors declare that the data supporting the findings of this study are available with the paper and its Supplementary information files. The data that support the findings of this study are available from the corresponding author upon reasonable request.

