## Supplemental Files for "Revisiting PFA-mediated tissue fixation chemistry: *FixEL* enables trapping of small molecules in the brain to visualize their distribution dynamics"

[2]ERATO (Exploratory Research for Advanced Technology, JST), Tokyo 102-0075, Japan.

[3]Department of Biomolecular Engineering, Graduate School of Engineering, Nagoya University, Nagoya 464-8603, Japan.

[4]Division of Structural Biology, University of Oxford, Oxford OX3 7BN, UK.

[5]Neurobiology Division, MRC Laboratory of Molecular Biology, Cambridge CB2 0QH, UK.

[6]Department of Neurophysiology, Keio University School of Medicine, Tokyo 160-8582, Japan.

‡ These authors contributed equally to this work.

***Corresponding Author:**

Prof. Itaru Hamachi

Department of Synthetic Chemistry & Biological Chemistry, Graduate School of Engineering, Kyoto University

### Supplementary Figures

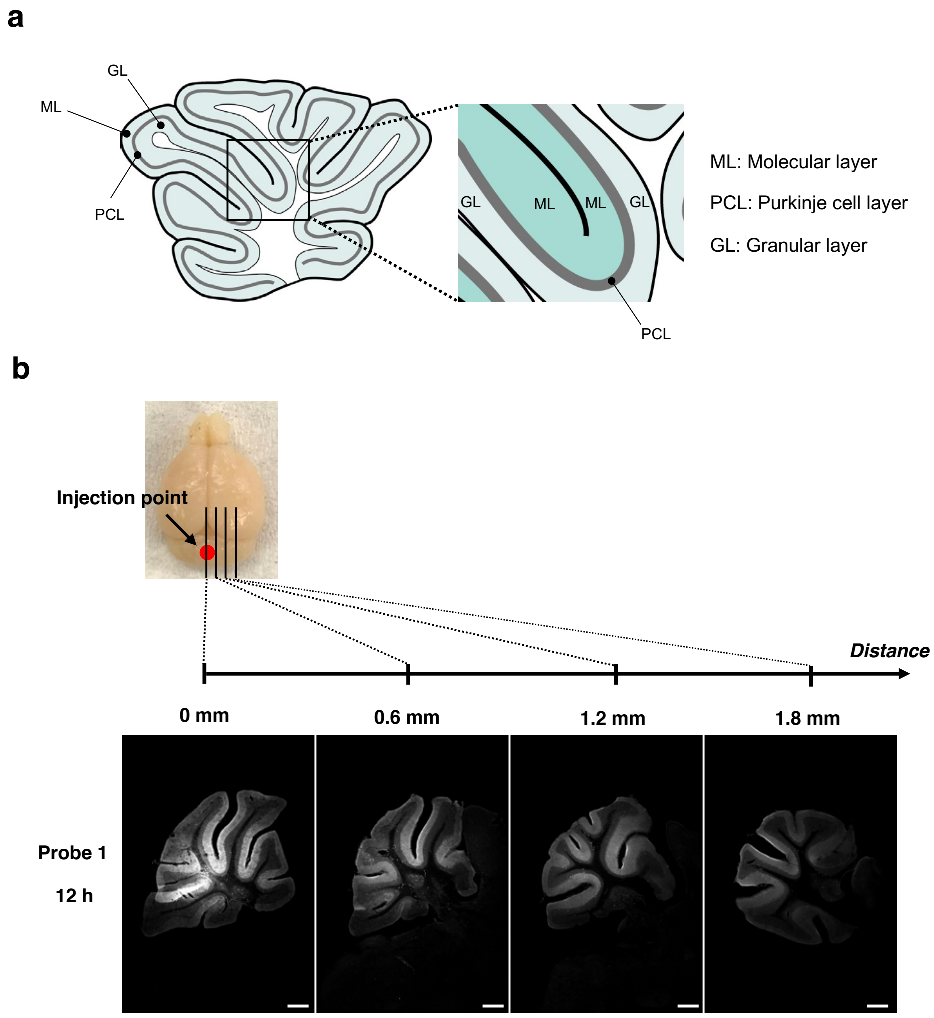


**Supplementary Figure 1 | Fluorescence imaging of cerebellum slices after *FixEL* with probe 1 at different distances from the midline of the cerebellum. a**, Sagittal slice image of the cerebellum. **b**, Fluorescence imaging of cerebellum slices after *FixEL* with probe **1**. PBS(–) containing 20 μM of probe **1** (4.5 μL) was injected into the mouse cerebellum. After 12 h of incubation, the mouse was transcardially perfused with 4% PFA. After slice preparation (40-μm thick), fluorescence imaging of the slices was performed using a CLSM equipped with a 5× objective and GaAsP detector (633 nm excitation for Ax647). Scale bar: 500 μm.

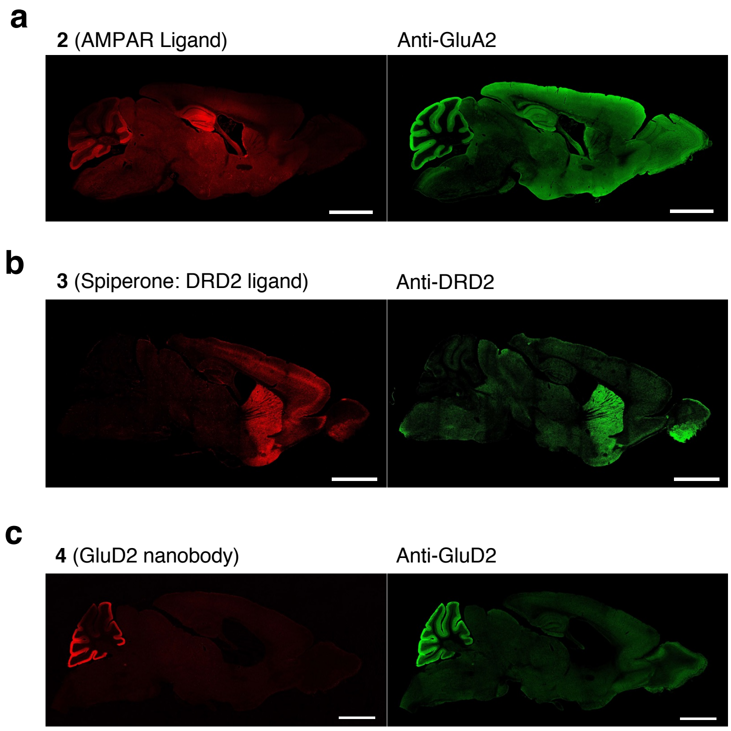


**Supplementary Figure 2 | Fluorescence imaging of whole-brain sagittal slices after *FixEL* using probe 2-4. a**, Co-immunostaining of whole-brain sagittal slices after *FixEL* with probe **2**. PBS(−) containing 40 μM of probe **2** (4.5 μL) was injected into the mouse lateral ventricle. After 3 h of incubation, mouse was transcardially perfused with 4% PFA. The slices (15-μm thick) after *FixEL* with probe **2** were treated with heat-induced-epitope-retrieval process with ImmunoSaver (FUJIFILM Wako: 80°C 20 min) and immunostained using anti-GluA2 (Merck, MAB397). Fluorescence imaging was performed using a CLSM equipped with a 10× objective and GaAsP detector (594 nm excitation for Ax594, 647 nm excitation for Ax647). Scale bar: 2 mm. **b**, Co-immunostaining of whole-brain sagittal slices after *FixEL* with probe **3**. PBS(−) containing 25 μM of probe **3** (4.5 μL) was injected into the mouse lateral ventricle. After 8 h of incubation, mouse was transcardially perfused with 4% PFA. The slices (50-μm thick) after *FixEL* with probe **3** were permeabilized and immunostained using anti-DRD2 (Frontier Institute, D2R-Rb-Af960). Fluorescence imaging was performed using a CLSM equipped with a 5× objective and GaAsP detector (488 nm excitation for Ax488, 633 nm excitation for Ax647). Scale bar: 2 mm. **c**, Co-immunostaining of whole-brain sagittal slices after *FixEL* with probe **4**. PBS(−) containing 10 μM of probe **4** (4.5 μL) was injected into the mouse lateral ventricle (left and right). After 24 h of incubation, mouse was transcardially perfused with 4% PFA. The slices (50-μm thick) after *FixEL* with probe **4** were permeabilized and immunostained using anti-GluD2 (SIGMA,

HPA056253). Fluorescence imaging was performed using a CLSM equipped with a 10× objective and GaAsP detector (488 nm excitation for Ax488, 633 nm excitation for Ax647). Scale bar: 2 mm.

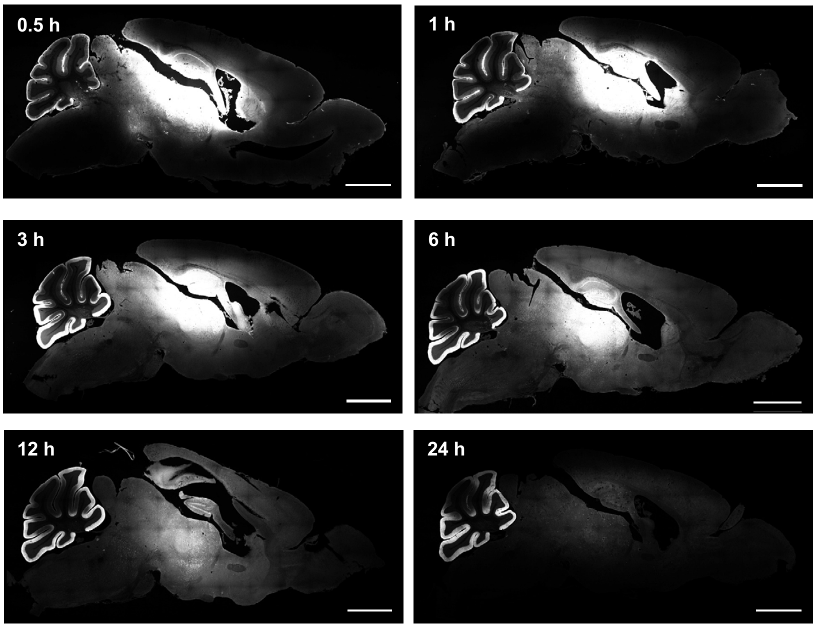


**Supplementary Figure 3 | Visualization of distribution dynamics on of probe 4 after *FixEL* at different incubation time.** PBS(−) containing 40 µM of probe **4** (4.5 µL) was injected into the mouse lateral ventricle. After 0.5, 1, 3, 6, 12, or 24 h of incubation, mouse was transcardially perfused with 4% PFA. Fluorescence imaging of slices (50-µm thick) was performed using a CLSM equipped with a 5× objective and GaAsP detector (633 nm excitation for Ax647). Scale bar: 2 mm.

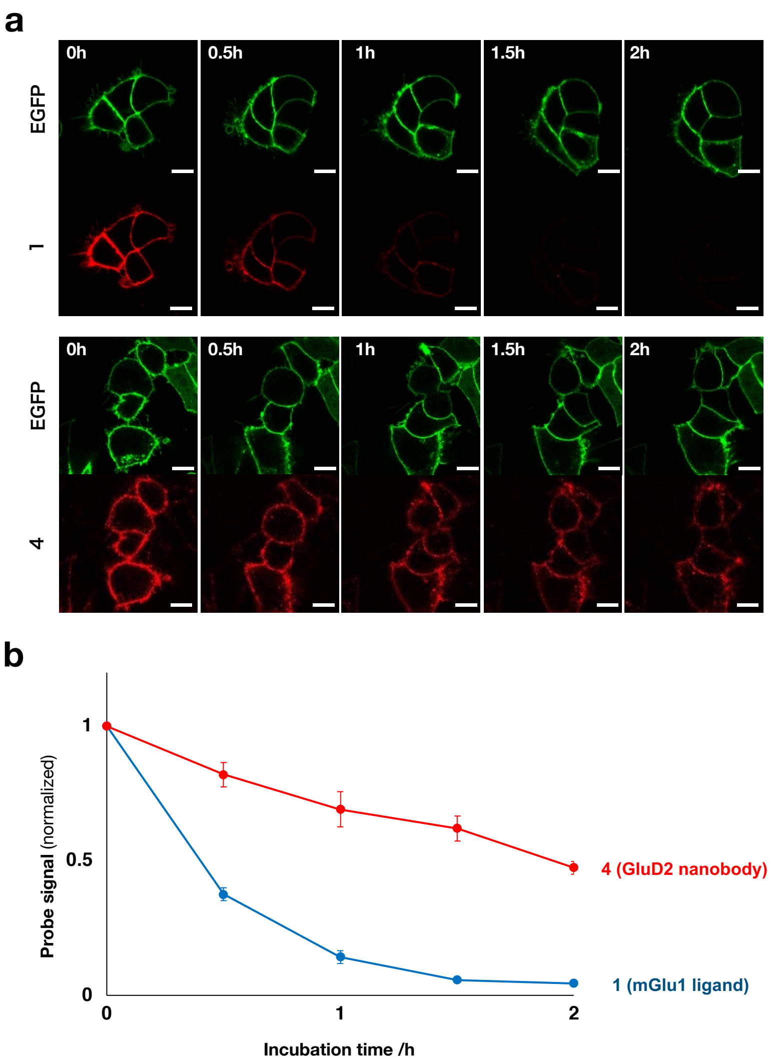


**Supplementary Figure 4 | Comparison of the dissociation rate between probe 1 and 4 in living cells. a,** Confocal imaging of mGlu1 or GluD2 and EGFP-F (green) expressed HEK293T cells after adding probe **1** or **4** (Ax647, red), washing with PBS(-), and incubation with DMEM in a 5% $CO_2$ humidified chamber at 37 °C for 0, 0.5, 1, 1.5, and 2h. Fluorescence imaging of the cells was performed using a CLSM equipped with a 40× objective and a GaAsP detector (488 nm excitation for EGFP and 633 nm excitation for Ax647). Scale bar 10 µm. **b,** Fluorescence change of the plasma membrane on mGlu1 or GluD2-expressed HEK293T cells depending on the incubation time with probe **1** (blue circle) or probe **4** (red square). n = 8 cells. Data are presented as mean ± s.e.m. Fluorescent signal of probe **4** from cell membrane still presented even after 2 hours of incubation, while probe **1** signal was almost disappeared after 1.5 hour of incubation.

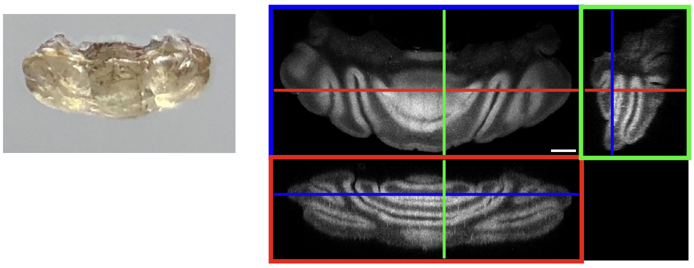

**Supplementary Figure 5 | 3D imaging of cerebellum regions after *FixEL* with probe 1 and 3DISCO.** PBS(–) containing 20 μM of probe **1** (4.5 μL) was injected into the mouse cerebellum. After 12 h of incubation, the mouse was transcardially perfused with 4% PFA. After 3DISCO treating, z-stacking fluorescence imaging of the cerebellum was performed using a CLSM equipped with a 5× objective and GaAsP detector (633 nm excitation for Ax647). Scale bar: 500 μm.

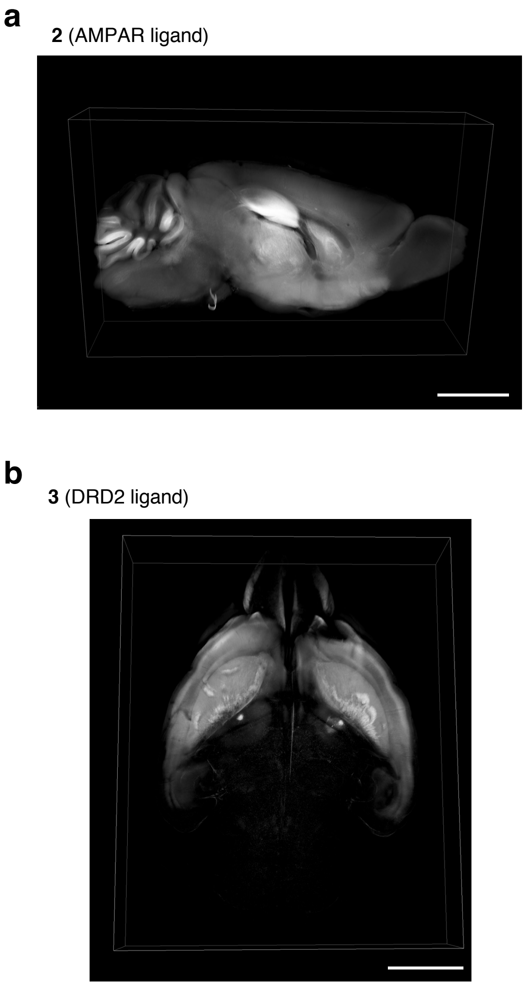


**Supplementary Figure 6 | 3D imaging of the brains after *FixEL* and 3DISCO tissue clearing.** **a**, Z-stacking fluorescence imaging of *FixEL* probe **2**. PBS(–) containing 40 μM of probe **2** (4.5 μL) was injected into mouse lateral ventricle. After 3 h of incubation, the mouse was transcardially perfused with 4% PFA. After 3DISCO treating, z-stacking fluorescence imaging of the whole brain was performed using a CLSM equipped with a 5× objective and GaAsP detector (633 nm excitation for Ax647). Scale bar: 2 mm. **b**, Z-stacking fluorescence imaging of *FixEL* probe **3**. PBS(–) containing 25 μM of probe **3** (4.5 μL) was injected into mouse lateral ventricle (left and right). After 20 h of incubation, the mouse was transcardially perfused with 4% PFA. After 3DISCO treating, z-stacking fluorescence imaging of the whole brain was performed using a CLSM equipped with a 5× objective and GaAsP detector (633 nm excitation for Ax647). Scale bar: 2 mm.

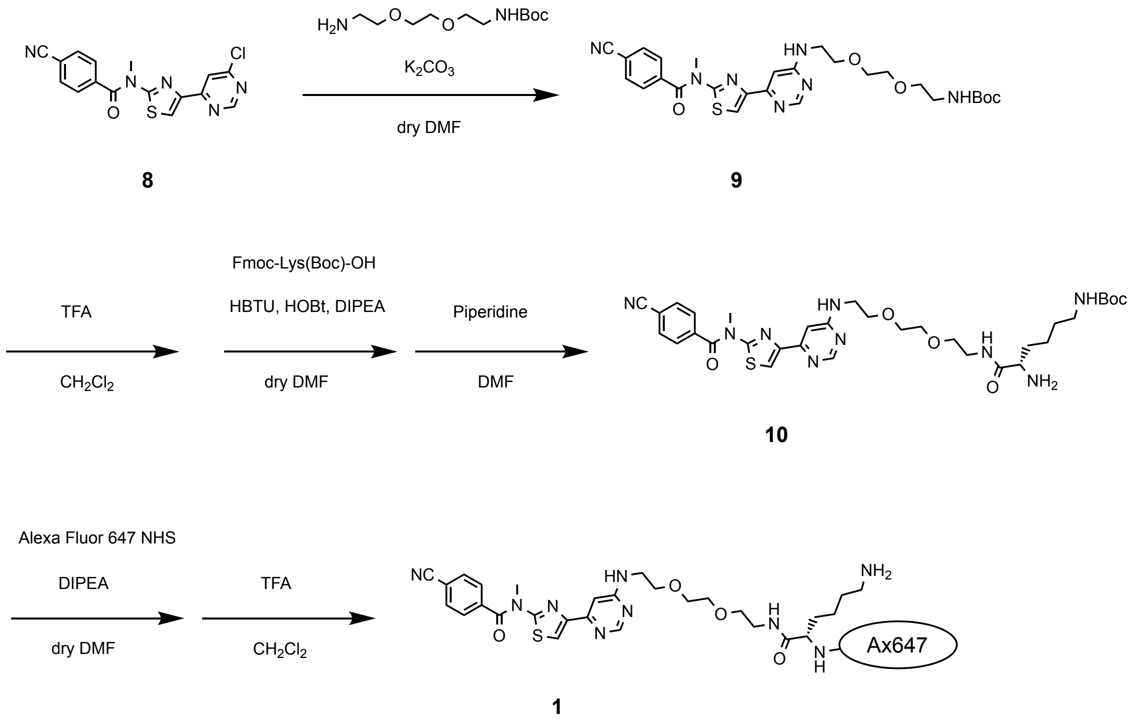


**Supplementary Figure 7 | Synthesis of probe 1.**

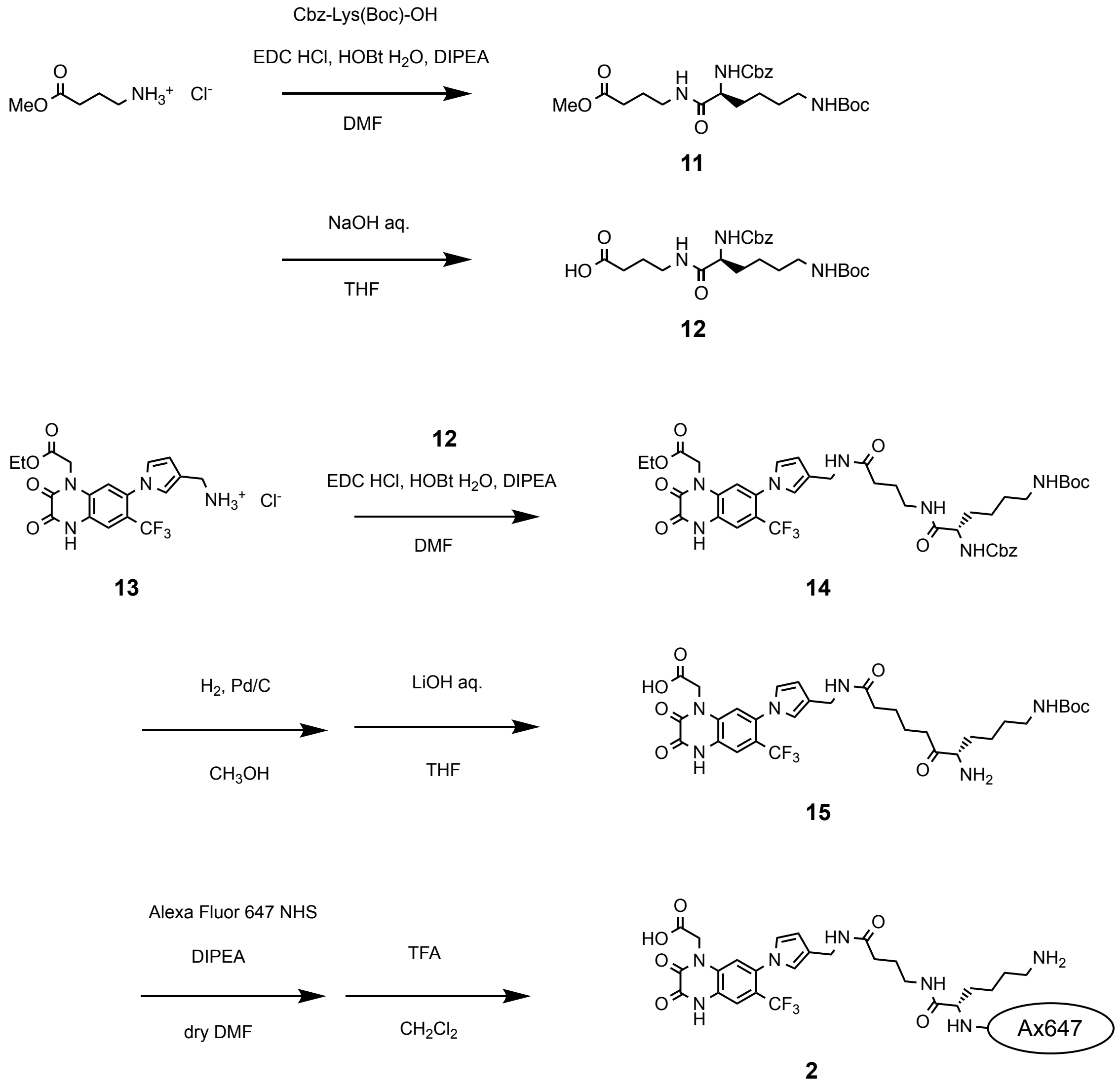


**Supplementary Figure 8 | Synthesis of probe 2.**

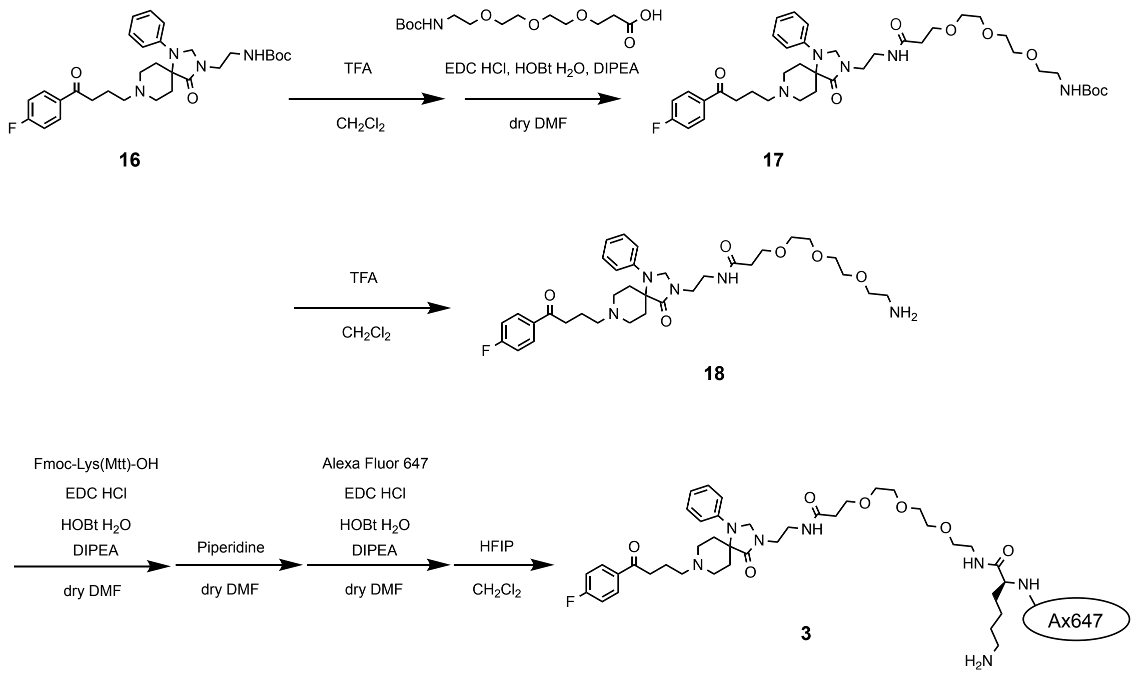


**Supplementary Figure 9 | Synthesis of probe 3.**

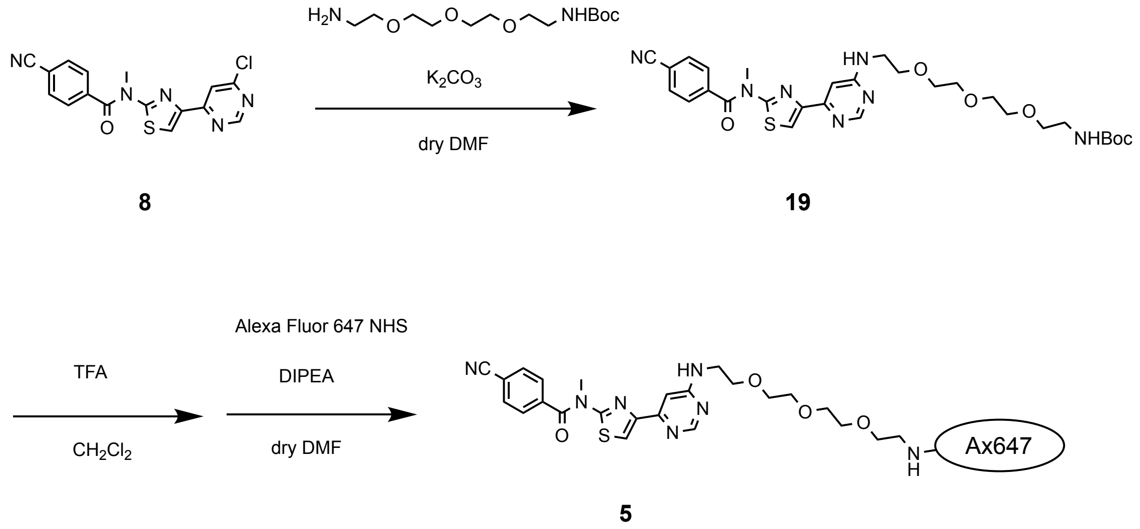


**Supplementary Figure 10 | Synthesis of probe 5.**

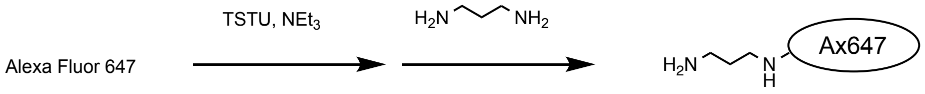


**6**

**Supplementary Figure 11 | Synthesis of probe 6.**

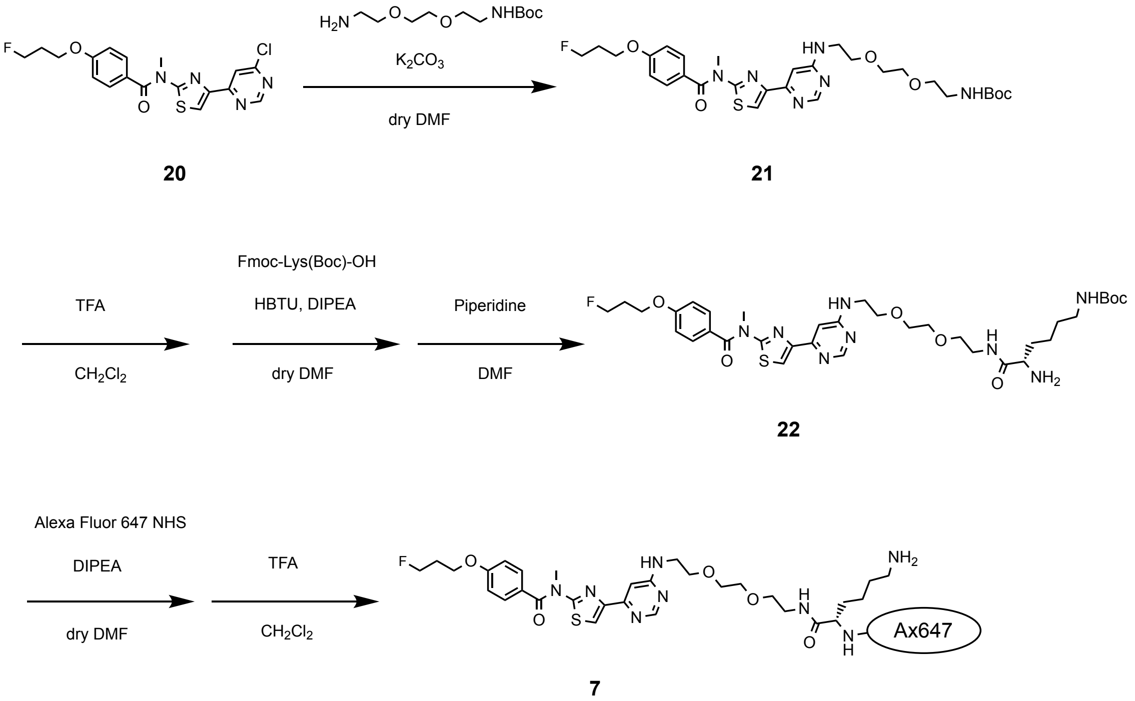


**Supplementary Figure 12 | Synthesis of probe 7.**

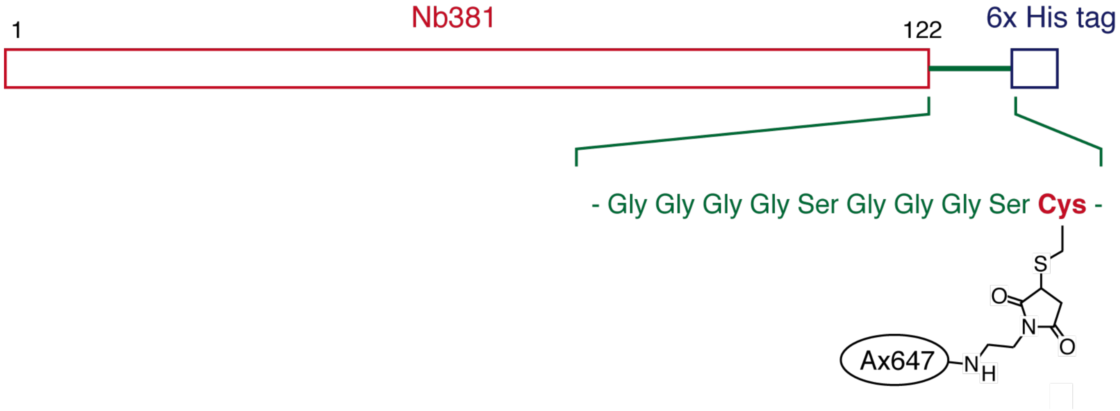


**Supplementary Figure 13 | Structure of Nb381-Ax647 (4).**

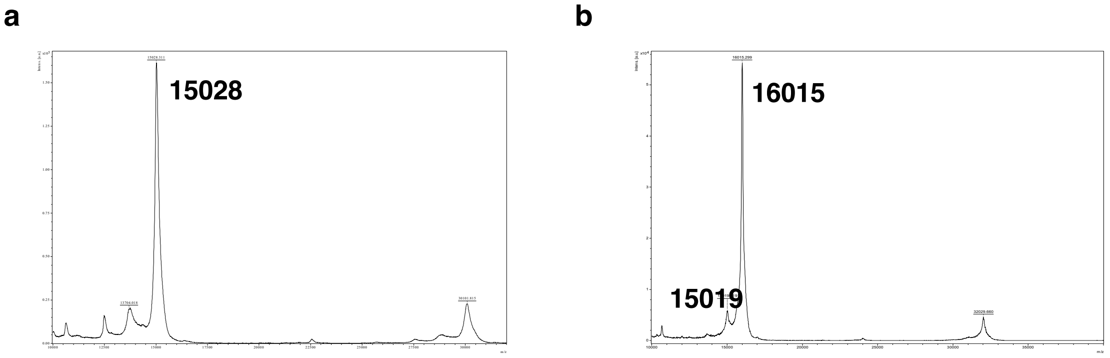


**Supplementary Figure 14 | MALDI TOF MS spectra of Nb381-(GS)2-Cys (a) and Nb381-Ax647 (b).**

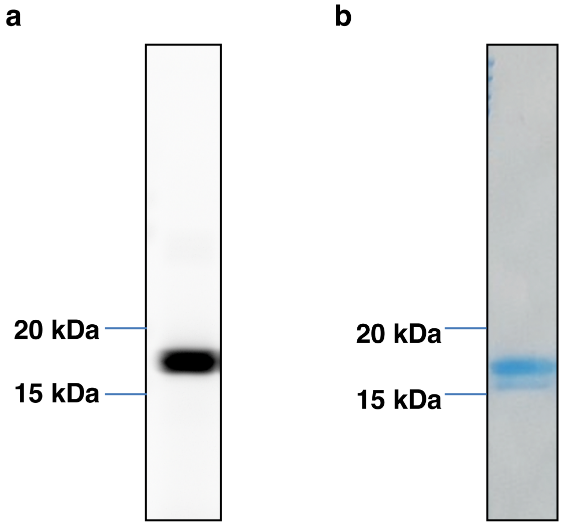


**Supplementary Figure 15 | SDS-PAGE analysis of Nb381-Ax647.** In Gel fluorescence (a) and CBB staining (b) images of Nb381-Ax647. 15% acryl amide gel. The modification yield was estimated to be 80% based on the band intensities of CBB-staining bands.

### Supplementary Methods

#### Synthesis and Characterization of Compounds

##### General materials and methods for organic synthesis

All chemical reagents and solvents were obtained from commercial suppliers (Aldrich, Tokyo Chemical Industry (TCI), Wako Pure Chemical Industries, or Watanabe Chemical Industries) and used without further purification. $^{1}$H-NMR spectra were recorded in deuterated solvents on a JEOL ECS400 or JEOL ECZ600R. Chemical shifts were referenced to residual solvent peaks or tetramethylsilane (δ = 0 ppm). Multiplicities are abbreviated as follows: s = singlet, d = doublet, t = triplet, m = multiplet. MALDI-TOF Mass spectra were measured on UltrafleXtreme (Bruker Daltonics). High resolution mass spectra were measured on an Exactive (Thermo Scientific) equipped with electron spray ionization (ESI). Reversed-phase HPLC (RP-HPLC) was carried out on a Hitachi Chromaster system equipped with a diode array.

##### Synthesis of 1

*tert*-butyl (2-(2-(2-((6-(2-(4-cyano-N-methylbenzamido)thiazol-4-yl)pyrimidin-4-yl)amino) ethoxy)ethoxy)ethyl)carbamate (**9**): The compound **8** was synthesized according to the literature.[S1] To a solution of compound **8** (16 mg, 45 μmol) in DMF (0.3 mL), *tert*-butyl (2-(2-(2-aminoethoxy)ethoxy)ethyl)carbamate (55.8 mg, 225 μmol, 5.0 eq.) and $K_2CO_3$ (37.5 mg, 270 μmol, 6.0 eq.) were added. The reaction mixture was stirred for 6.5 h at 50 °C. The solvent was evaporated and the residue was purified by silica gel column chromatography ($CHCl_3$:MeOH = 9:1) to yield compound **9** (11 mg, 19 μmol, 43%) as a yellow oil. $^{1}$H NMR (400 MHz, $CDCl_3$): δ 8.57 (s, 1H), 7.94 (s, 1H), 7.80 (d, $J$ = 8.4 Hz, 2H), 7.67 (d, $J$ = 8.4 Hz, 2H), 7.07 (s, 1H), 3.71–3.63 (m, 11H), 3.55 (t, $J$ = 5.2 Hz, 2H), 3.33 (m, 2H), 1.41 (s, 9H). $^{13}$C NMR (150 MHz, $CDCl_3$): δ 168.43, 163.30, 159.90, 158.61, 157.31, 156.13, 148.35, 138.71, 132.61, 128.23, 117.71, 116.30, 114.91, 100.18, 79.34, 70.53-69.62, 41.06, 40.53, 38.13, 28.48. HR-ESI-MS: calcd. for $C_{27}H_{33}N_7O_5S$ $[M+Na]^+$ = 590.2156, obsd = 590.2150.

*tert*-butyl (*S*)-(5-amino-6-((2-(2-(2-((6-(2-(4-cyano-N-methylbenzamido)thiazol-4-yl)pyrimidin-4-yl)amino)ethoxy)ethoxy)ethyl)amino)-6-oxohexyl)carbamate (**10**): To a stirred solution of compound **9** (3.0 mg, 6.4 μmol) in $CH_2Cl_2$ (0.5 mL), trifluoroacetic acid (TFA) (0.5 mL) was added and the reaction solution was stirred at RT for 30 min. After azeotropic removal of TFA with toluene (2 mL x 2), the residue was dissolved in dry DMF (100 μL). Then, Fmoc-Lys(Boc)-OH (4.5 mg, 9.6 μmol, 1.5 eq.), HBTU (7.3 mg, 19 μmol, 3.0 eq.), HOBt (3.0 mg, 20 μmol, 3.1 eq.) and DIPEA (2.6 μL, 58 μmol, 9.0 eq.) were added. After stirring the reaction mixture at RT for 2 h, piperidine (10 μL) was added and the solution was stirred at RT for another 10 min. The solvent was evaporated and the residue was purified by reversed-phase HPLC with a linear gradient of 30-60 % $CH_3CN$-0.1% TFA (30 min). After lyophilization, the compound **10** was obtained as a yellow oil (4.8 mg, 92%). $^1$H NMR (400 MHz, $CD_3OD$): δ 8.58 (s, 1H), 8.13 (s, 1H), 7.84 (d, $J$ = 8.0 Hz, 2H), 7.72 (d, $J$ = 8.0 Hz, 2H), 7.20 (s, 1H), 3.73 (s, 3H), 3.63–3.57 (m, 11H), 3.48 (m, 2H), 2.93 (t, $J$ = 6.0 Hz, 2H), 1.43–1.32 (m, 15H). $^{13}$C NMR (150 MHz, $CDCl_3$): δ170.84, 170.21, 165.21, 162.53, 158.63, 154.16, 148.91, 141.87, 139.66, 133.81, 129.56, 120.32, 118.83, 115.95, 102.30, 80.08, 71.53, 71.32, 70.39, 70.10, 54.50, 42.66, 40.88, 40.52, 38.74, 32.33, 30.57, 28.80, 23.08. HR-ESI-MS: calcd. For $C_{33}H_{45}N_9O_6S$ $[M+H]^+$ = 696.3286, obsd. = 696.3281.

Compound **1**: To a solution of compound **10** (0.4 mg, 0.5 μmol) in dry DMF (50 μL), Alexa Fluor™ 647 NHS ester (1.0 mg, 0.8 μmol, 1.6 eq.) and DIPEA (1 μL, 5.7 μmol, 11 eq.) were added and the reaction mixture was stirred at RT for 1 h. The reaction mixture was purified by reversed-phase HPLC with a linear gradient of 10-40 % $CH_3CN$-0.1% TFA (30 min). After azeotropic removal of TFA with toluene (2 mL x 2), the residue was dissolved in $CH_2Cl_2$/TFA (1/1) (1 mL) and stirred at RT for 30 min. After azeotropic removal of TFA with toluene (2 mL x 2), the solvent was removed to afford **1** as a blue oil (0.32 mg, 44%). HR-ESI-MS: calcd. for $C_{64}H_{81}N_{11}O_{17}S_5$ $[M-3H+2Na]^-$ = 1478.3982, obsd. = 1478.4004. calcd. for $[M-2H+Na]^-$ = 1456.4162, obsd. = 1456.4191.

**Synthesis of 2**

Methyl (*S*)-4-(2-(((benzyloxy)carbonyl)amino)-6-((*tert*-butoxycarbonyl)amino)hexanamido)butanoate (**11**): To a solution of *N*-*α*-benzyloxycarbonyl-*N*-*ε*-*tert*-butoxycarbonyl-*L*-lysine (500 mg, 1.31 μmol, 1.0 eq.) in dry DMF (13 mL) were added 4-aminobutyric acid methyl ester hydrochloride (240 mg, 1.58 μmol, 1.2 eq.), 1-(3-dimethylaminopropyl)-3-ethylcarbodiimide hydrochloride (EDC·HCl: 328 mg, 1.71 μmol, 1.3 eq.), 1-hydroxybenzotriazole monohydrate (HOBt·$H_2O$: 262 mg, 1.71 μmol, 1.3 eq.), and DIPEA (916 μL, 5.26 μmol, 4.0 eq.). After stirring for 10 h, the reaction mixture was diluted with EtOAc (60 mL) and then washed with 0.01 M HCl aq. (20 mL x2), saturated $NaHCO_3$ aq. (20 mL x2) and brine (20 mL). The organic layer was dried over $Na_2SO_4$, and evaporated under reduced pressure. The residue was purified by silica gel column chromatography ($CHCl_3$:MeOH:AcOH = 100:2:0.5 to 100:5:0.5) to yield compound **11** (613 mg, 97%) as a white solid. $^1H$ NMR (400 MHz, $CD_3OD$): δ 7.36–7.26 (m, 5H), 5.07 (s, 2H), 4.02–3.98 (m, 5H), 3.64 (s, 3H), 3.31–3.29 (m, 2H), 3.03–2.98 (m, 2H), 2.36–2.32 (m, 2H), 1.81–1.72 (m, 2H), 1.69–1.59 (m, 2H), 1.41 (s, 9H), 1.37–1.31, (m, 4H). $^{13}C$ NMR (100 MHz, $CD_3OD$): δ 175.28, 175.03, 158.52, 158.38, 138.13, 129.46, 129.01, 128.90, 128.78, 79.82, 67.67, 56.54, 52.07, 40.95, 39.53, 32.93, 31.91, 30.53, 28.79, 25.64, 24.16.

(*S*)-4-(2-(((Benzyloxy)carbonyl)amino)-6-((*tert*-butoxycarbonyl)amino)hexanamido)butanoic acid **(12):** To a solution of **11** (599 mg, 1.25 mmol) in THF (12.5 mL) was added portionwise 1 M NaOH (1.25 mL, 1.25 mmol, 1.0 eq.) every 1 h for 4 h. After stirring for another 1 h, the pH of reaction mixture was adjusted to pH 3 using 1 M HCl aq. The crude product was extracted with EtOAc (60 mLx1 and 30 mLx2). The organic layer was washed with water (20 mL x 2) and brine (20 mL x 1). The organic layer was dried over $Na_2SO_4$ and evaporated under reduced pressure. The residue was purified by silica gel column chromatography ($CHCl_3$:MeOH:AcOH = 100:2:0.5) to afford **12** (393 mg, 68%) as a white solid. $^1H$ NMR (400 MHz, $CD_3OD$) δ 7.34–7.27 (m, 5H), 5.07 (s, 2H), 4.04–4.01 (m, 1H), 3.21 (t, $J$ = 6.4 Hz, 2H), 3.00 (t, $J$ = 6.4 Hz, 2H), 2.33–2.29 (m, 2H), 1.79–1.62 (m, 4H), 1.41 (s, 9H), 1.36–1.32 (m, 4H). $^{13}C$ NMR (100 MHz, $CD_3OD$): δ 176.84, 175.16, 175.01, 158.51, 158.38, 138.09, 129.91, 129.45, 129.20, 129.01, 126.29, 79.83, 67.69, 56.61, 40.94, 39.66, 32.93, 32.07, 30.51, 28.78, 25.67, 24.14.

Ethyl (*S*)-2-(7-(3-(9-(((benzyloxy)carbonyl)amino)-17,17-dimethyl-3,8,15-trioxo-16-oxa-2,7,14-triazaoctadecyl)-1*H*-pyrrol-1-yl)-2,3-dioxo-6-(trifluoromethyl)-3,4-dihydroquinoxalin-1(2*H*)-yl)acetate (**14**): The compound **13** was synthesized according to the literature.[S2,S3] To a solution of **13** (30 mg, 67 μmol, 1.0 eq.) in dry DMF (2.0 mL) were added **12** (40.3 mg, 87 μmol, 1.3 eq.), 1-(3-dimethylaminopropyl)-3-ethylcarbodiimide hydrochloride (EDC·HCl: 16.7 mg, 87 μmol, 1.3 eq.), 1-hydroxybenzotriazole monohydrate (HOBt·$H_2O$: 13,4 mg, 87 μmol, 1.3 eq.), and DIPEA (47 μL, 269 μmol, 4.0 eq.) under $N_2$ atmosphere. After stirring for 17 h, the resulting mixture was evaporated under reduced pressure. The resulting residue was dissolved in

$CHCl_3$ (100 mL) and washed with 0.01 M HCl aq. (20 mL x 2), saturated $NaHCO_3$ (20 mL x 2) and brine (20 mL). The organic layer was dried over $Na_2SO_4$ and evaporated under reduced pressure. The residue was purified by silica gel column chromatography ($CHCl_3$:MeOH = 100:5 to 100:10) to afford **14** as a white solid (47.6 mg, 91%). $^1$H NMR (400 MHz, $CD_3OD$) δ 7.59 (s, 1H), 7.33–7.19 (m, 6H), 6.81–6.70 (m, 2H), 6.21–6.19 (m, 1H), 5.05–4.95 (m, 4H), 4.24 (s, 2H), 4.20 (q, *J* = 7.2 Hz, 2H), 4.02–3.99 (m, 1H), 3.25–3.18 (m, 2H), 3.03–2.95 (m, 2H), 2.20 (t, *J* =7.2 Hz, 2H), 1.81–1.55 (m, 4H), 1.43 (s, 9H), 1.36–1.28 (m, 4H), 1.25 (t, *J* = 7.2 Hz, 3H). $^{13}$C NMR (100 MHz, $CD_3OD$): δ 174.70, 174.62, 175.01, 168.22, 158.12, , 157.97, 156.79, 154.62, 137.43, 135.91, 130.49, 129.17, 128.80, 128.60, 125.92, 125.07, 124.81, 123.19, 123.03, 122.87, 122.57, 122.36, 117.08, 115.72, 115.67, 110.29, 79.69, 67.53, 63.08, 58.11, 56.33, 45.37, 40.65, 39.42, 33.98, 32.62, 30.19, 28.75, 26.40, 23.79, 18.25, 14.38.

(*S*)-2-(7-(3-(9-amino-17,17-dimethyl-3,8,15-trioxo-16-oxa-2,7,14-triazaoctadecyl)-1*H*-pyrrol-1-yl)-2,3-dioxo-6-(trifluoromethyl)-3,4-dihydroquinoxalin-1(2*H*)-yl)acetic acid (**15**): To a solution of **14** (44.4 mg, 51.8 μmol) in dry MeOH (2.0 mL) was added 10% Pd/C (20 mg) under $N_2$ atmosphere. After stirring for 4 h under $H_2$ atmosphere, the resulting mixture was filtered with a Celite® pad and evaporated under reduced pressure to afford the Cbz deprotected product. The Cbz deprotected product was dissolved in THF (2 mL) and 0.5 M LiOH aq. (0.31 mL, 155 μmol, 3.0 eq.) was added. After stirring for 2 h, the resulting mixture was neutralized with 1 M HCl aq. and evaporated under reduced pressure. The crude product was dissolved in $CHCl_3$ : MeOH = 3 :1 (2 mL ), then filtered. The filtrate was evaporated under reduced and the residue was purified by silica gel column chromatography ($CHCl_3$:MeOH:28% NH3 aq. = 100:50:3 to 100:100:4) to afford **15** as a white solid (23.7 mg, 66%). $^1$H NMR (400 MHz, $CD_3OD$) δ 7.57 (s, 1H), 7.14 (s, 1H), 6.80 (bs, 1H), 6.76 (bs, 1H), 6.21–6.17 (m, 1H), 4.76 (s, 2H), 4.27, (d, *J* = 14.8 Hz, 1H), 4.22 (d, *J* = 14.8 Hz, 1H), 3.75 (t, *J* = 6.8 Hz, 1H), 3.26–3.17 (m, 2H), 3.02 (t, *J* = 7.2 Hz, 2H), 2.25 (t, *J* = 7.2 Hz, 2H), 1.88–1.69 (m, 4H), 1.53–1.32 (m, 4H), 1.41 (s, 9H). $^{13}$C NMR (100 MHz, $CD_3OD$): δ 174.90, 173.23, 171.20, 158.59, 157.55, 135.93, 131.65, 125.09, 123.53, 122.89, 117.75, 110.49, 79.94, 58.33, 54.68, 40.89, 39.96, 37.18, 34.34, 32.73, 30.54, 28.80, 27.22, 23.34, 18.38.

Compound **2**: To a solution of **15** (1.0 mg, 1.46 μmol, 1.4 eq.) in dry DMF (0.1 mL) was added Alexa Fluor™ 647 NHS ester (1.0 mg, 1.04 μmol, 1.0 eq.) and DIPEA (1.09 μL, 3.13 μmol, 6.0 eq.) under $N_2$ atmosphere. After stirring for 4 h, the resulting mixture was evaporated under reduced pressure. The crude product was dissolved in $CH_2Cl_2$ (1.0 mL). To this solution was added TFA (200 μL). After stirring for 1 h, the solvent was removed azeotropically with toluene (200 μL, x3). The crude product was purified by RP-HPLC (YMC-Pack ODS-A, $CH_3CN$ : $CH_3COONH_4$ = 15 : 85 → 30 : 70, linear gradient over 45 min) to afford **2**. HR-ESI-MS: calcd. for $C_{76}H_{104}FN_9O_{20}S_4$ $[M-2H]^{2-}$ = 716.6977, obsd. = 716.6962.

**Synthesis of 3**

*tert*-butyl (15-(8-(4-(4-fluorophenyl)-4-oxobutyl)-4-oxo-1-phenyl-1,3,8-triazaspiro[4.5]decan-3-yl)-12-oxo-3,6,9-trioxa-13-azapentadecyl)carbamate (**17**): The compound **16** was synthesized according to the literature.[S4] To a solution of **16** (122 mg, 0.23 mmol) in $CH_2Cl_2$ (4 mL) were added TFA (1 mL). After stirring for 1 h at RT under Ar atmosphere, the solvent was removed azeotropically with toluene (1 mL x 3) and $CHCl_3$ (1 mL x 3) to afford the Boc deprotected product. The Boc deprotected product (80 mg, 0.18 mmol, 1 eq.) was dissolved in dry DMF (10 mL), and Boc- 3-(2-(2-(2-aminoethoxy)ethoxy)ethoxy)propanoic acid (76.2 mg, 0.25 mmol, 1.4 eq.), EDC HCl (52.5 mg, 0.27 mol, 1.5 eq.), HOBt $H_2O$ (41.9 mg, 0.25 mmol, 1.5 eq) and DIPEA (283 μL, 1.8 mmol, 10 eq.) were added. After stirring at RT for 16 h, $Et_2O$ (40 mL) was added and the organic layer was washed with sat. $NaHCO_3$ aq. (15 mL x 3). The organic layer was dried over $MgSO_4$ and evaporated under reduced pressure to afford compound **17** (72 mg, 0.095 mmol, 53%) as a white solid. $^1H$ NMR (400 MHz, $CDCl_3$) δ 8.02 (dd, *J* = 7.2 Hz, 5.6 Hz, 2H), 7.24 (t, *J* = 8.4 Hz, 2H), 7.13 (t, *J* = 8.4 Hz, 2H), 6.90 – 6.85 (m, 3H), 5.16 (br, 1H), 4.74 (s, 2H), 3.71 (t, *J* = 6.0 Hz, 2H), 3.64 – 3.50 (m, 15 H), 3.31 – 3.20 (m, 2H), 3.01 (t, *J* = 7.2 Hz, 2H), 2.81 – 2.79 (m, 4H), 2.60 – 2.44 (m, 6H), 1.96 (t, *J* = 7.2 Hz, 2H), 1.44 (s, 9H), 0.89 (t, *J* = 6.4 Hz, 2H)

3-(2-(2-(2-aminoethoxy)ethoxy)ethoxy)-*N*-(2-(8-(4-(4-fluorophenyl)-4-oxobutyl)-4-oxo-1-phenyl-1,3,8-triazaspiro[4.5]decan-3-yl)ethyl)propanamide (**18**): To a solution of **17** (72 mg, 95 μmol) in $CH_2Cl_2$ (9 mL) were added TFA (1 mL). After stirring for 1 h at RT under Ar atmosphere, the solvent was removed azeotropically with toluene (6 mL x 1) and $CH_2Cl_2$ (3 mL x 3). The residue was purified by RP-HPLC (Cosmosil 5C18AR2, $CH_3CN$-0.1% TFA : $H_2O$-0.1% TFA = 20 : 80 → 70 : 30, linear gradient over 50 min) to afford **18** (8.0 mg, 9% as 3TFA salts) as a yellow oil. $^1H$ NMR (400 MHz, $CD_3OD$) δ 8.08 (dd, *J* = 8.8 Hz, 5.6 Hz, 2H), 7.30 (t, *J* = 8.4 Hz, 2H), 7.23 (t, *J* = 8.4 Hz, 2H), 7.02 (d, J = 8.8 Hz, 2H), 6.93 (t, *J* = 5.6 Hz, 1H), 4.84 (s, 2H), 3.82 (ddd, *J* =13.2 Hz, 2.8 Hz, 2H ) 3.70 – 3.48 (m, 18H), 3.40 (t , *J* = 5.6 Hz, 2H), 3.26 – 3.19 (m, 4H), 3.01 (t, 4.8 Hz, 2H), 2.79 (br, 2H),2.44 (t, *J* = 6.0 Hz, 2H), 2.16(br, 2H), 2.04 (br, 2H)

Compound **3**: Compound **18** (8 mg, 12.5 μmol, 1 eq.) was dissolved in dry DMF (2 mL), and Fmoc-Lys(Mtt)-OH (10.5 mg, 16.8 μmol, 1.35 eq.), EDC HCl (4.2 mg, 21.9 μmol, 1.8 eq.), HOBt $H_2O$ (2.5 mg, 16.3 μmol, 1.31 eq.), and DIPEA (25.7 μL, 165.7 μmol, 13.3 eq.) were added. After stirring at RT for 18 h, $CH_2Cl_2$ (10 mL) was added and the organic layer was washed with sat. $NaHCO_3$ aq. (2 mL x 2). The organic layer was dried over $Na_2SO_4$ and evaporated under reduced pressure. The residue was dissolved with dry DMF (0.8 mL) and piperidine (0.2 mL). The mixture was stirred at RT for 90 min and the solvent was removed by evaporation under reduced pressure to give a Fmoc deprotected product. The Fmoc deprotected product (1.7 mg, 1.66 μmol, 1 eq.) was dissolved in dry DMF (1 mL) and Alexa Fluor™ 647 (1.9 mg, 2.21 μmol, 1.33 eq.), EDC HCl (0.49 mg, 2.56 μmol, 1.5 eq.), HOBt $H_2O$ (0.38 mg, 2.48 μmol, 1.5 eq.) and DIPEA (6 μL, 38.7 μmol, 23.4 eq.) were added under Ar atmosphere. After stirring at RT for 15 h, the solvent was removed by evaporation under reduced pressure. The crude product was dissolved in $CH_2Cl_2$ (0.54 mL) and HFIP (0.13 mL) was added. After stirring for 1 h, the solvent was removed and the residue was purified by RP-HPLC (Cosmosil 5C18AR2, $CH_3CN$-0.1% TFA : $H_2O$-0.1% TFA = 5 : 95 → 55 : 45, linear gradient over 50 min) to afford **3** as a blue film. HR-ESI-MS: calcd. for $C_{76}H_{104}FN_9O_{20}S_4$ $[M-3H+Na]^{2-}$ = 803.8049, obsd. =803.8060.

**Synthesis of 5**

*tert*-butyl (2-(2-(2-(2-((6-(2-(4-cyano-*N*-methylbenzamido)thiazol-4-yl)pyrimidin-4-yl)amino)ethoxy)ethoxy)ethoxy)ethyl)carbamate (**19**): To a solution of **8** (7.5 mg, 21 μmol) in DMF (0.1 mL) was added *tert*-butyl (2-(2-(2-(2-aminoethoxy)ethoxy)ethoxy)ethyl)carbamate (31 mg, 105 μmol, 5.0 eq.) and $K_2CO_3$ (15.0 mg, 105 μmol, 5.0 eq.). The resulting mixture was stirred at 50 °C. After completion of the reaction (22.5 h), DMF was removed in vacuo. The resulting residue was purified with flash column chromatography ($CH_2Cl_2$:MeOH = 9:1) to afford the compound **19** as yellow oil (4.5 mg, 35%). $^1$H NMR (600 MHz, $CDCl_3$): δ 8.58 (s, 1H), 7.96 (s, 1H), 7.83 (d, $J$ = 4.0 Hz, 2H), 7.70 (d, $J$ = 4.0 Hz, 2H), 7.11 (s, 1H), 3.73–3.63 (m, 15H), 3.55 (t, $J$ = 5.2 Hz, 2H), 3.31 (m, 2H), 1.42 (s, 9H). $^{13}$C NMR (150 MHz, $CDCl_3$): δ 168.43, 163.34, 159.89, 158.61, 157.08, 156.05, 148.34, 138.71, 132.61, 128.23, 117.71, 116.25, 114.90, 100.44, 77.03, 70.63-69.69, 40.99, 40.52, 38.12, 28.47. HR-ESI-MS: calcd. for $C_{29}H_{37}N7O_6S$ $[M+Na]^+$ = 634.2418, obsd. = 634.2410.

Compound **5**: To a solution of compound **19** (0.4 mg, 0.65 μmol, 1.0 eq.) in $CH_2Cl_2$ (0.5 mL), TFA (0.5 mL) was added and the reaction mixture was stirred at RT for 30 min. After azeotropic removal of TFA with toluene (2 mL x 2), the residue was dissolved in dry DMF (150 μL). To the solution, Alexa Fluor™ 647 NHS ester (1.0 mg, 0.8 μmol, 1.2 eq.) and DIPEA (1 μL, 6.3 μmol, 10 eq.) were added and the reaction mixture was stirred at RT for 1 h. The reaction mixture was purified by reversed-phase HPLC with a linear gradient of 20-40 % $CH_3CN$-0.1% TFA (20 min) to afford **5** as blue oil (0.2 mg, 24%). HR-ESI-MS: calcd. for $C_{60}H_{73}N_9O_{17}S_5$ $[M-H]^-$ = 1350.3655, obsd. = 1350.3677.

**Synthesis of 6**

Compound **6:** To a solution of Alexa Fluor 647 (0.5 mg, 0.5 μmol) in dry DMF (50 μL), *N,N,N',N'*-Tetramethyl-*O*-(*N*-succinimidyl)uronium tetrafluoroborate (TSTU: 0.9 mg, 2.9 μmol, 6 eq.) and triethylamine (3.2 μL, 23 μmol, 40 eq.) were added. After stirring at RT for 1 h, propane-1,3-diamine hydrochloride (0.9 mg, 5.8 μmol, 10 eq.) was added. After stirring at RT for 30 min, the reaction mixture was diluted with water and purified by reversed-phase HPLC with a linear gradient of 10-40 % $CH_3CN$-0.1% TFA (30 min) to afford **6**. HR-ESI-MS: calcd. for $C_{39}H_{54}N_4O_{13}S_4$ $[M-3H+Na]^{2-}$ = 467.1122, obsd. = 467.1123.

**Synthesis of 7**

*tert*-butyl (2-(2-(2-((6-(2-(4-(3-fluoropropoxy)-*N*-methylbenzamido)thiazol-4-yl)pyrimidin-4-yl)amino)ethoxy)ethoxy)ethyl)carbamate (**21**): The compound **20** was synthesized according to the literature.[S5] To a solution of compound **20** (108 mg, 0.265 mmol) in DMF (1 mL), *tert*-butyl (2-(2-(2-aminoethoxy)ethoxy)ethyl)carbamate (85 mg, 0.345 mmol, 1.3 eq.) and $K_2CO_3$ (55 mg, 0.398 mmol, 1.5 eq.) were added. After stirring at 60 °C overnight under Ar atmosphere, $CH_2Cl_2$ (50 mL) was added and the organic layer was washed with brine (25 mL x2). The organic layer was dried over $Na_2SO_4$ and the solvent was evaporated. The residue was purified by silica gel column chromatography ($CH_2Cl_2$ : CH3OH = 96 : 4) to yield compound **21** as a white solid (132 mg, 0.214 mmol, 81%). $^1H$ NMR (400 MHz, $CDCl_3$): δ 8.61 (d, *J* = 1.1 Hz, 1H), 7.93 (s, 1H), 7.60 (d, *J* = 8.8 Hz, 2H), 7.14 (s, 1H), 7.02 (d, *J* = 8.8 Hz, 2H), 4.69 (dt, *J* = 47.0, 5.7 Hz 2H), 4.20 (t, *J* = 6.1 Hz, 2H), 3.80 (s, 3H), 3.77–3.74 (m, 2H), 3.69–3.66 (m, 6H), 3.39 (t, *J* = 5.1 Hz, 2H), 3.38–3.37 (m, 2H), 2.32–2.15 (m, 2H), 1.46 (s, 9H). $^{13}C$ NMR (150 MHz, $CDCl_3$, 60℃): δ 170.04, 163.28, 160.98, 160.80, 158.39, 157.40, 155.98, 148.06, 129.81, 126.65, 115.58, 114.44, 99.91, 80.31, 79.09, 70.36, 70.29, 70.23, 69.58, 63.92, 40.96, 40.56, 38.42, 30.27, 28.33.

*tert*-butyl (*S*)-(5-amino-6-((2-(2-(2-((6-(2-(4-(3-fluoropropoxy)-*N*-methylbenzamido)thiazol-4-yl)pyrimidin-4-yl)amino)ethoxy)ethoxy)ethyl)amino)-6-oxohexyl)carbamate (**22**): To a solution of compound **21** (40 mg, 65 μmol) in $CH_2Cl_2$ (0.5 mL), TFA (0.5 mL) was added and the reaction solution was stirred at RT for 30 min. After azeotropic removal of TFA with toluene (2 mL x 2), the residue was dissolved in dry DMF (0.65 mL). Then, Fmoc-Lys(Boc)-OH (45 mg, 97 μmol, 1.5 eq.), HBTU (37 mg, 97 μmol, 1.5 eq.) and DIPEA (33 μL, 190 μmol, 2.9 eq.) were added. After stirring the reaction mixture at RT overnight, piperidine (98 μL) was added and the reaction mixture was stirred at RT for another 2 h. The solvent was evaporated and the residue was dissolved in $CH_2Cl_2$ (30 mL). The organic layer was washed with sat. $NaHCO_3$ aq. (15 mL x 2) and brine (15 mL x2). After drying over $Na_2SO_4$, the solvent was evaporated and the residue was purified by preparative TLC ($CH_2Cl_2$ : MeOH : NH3 aq. = 150 : 10 : 1.6) to yield compound **22** as a pale yellow form (46 mg, 60 μmol, 93%). $^1H$ NMR (400 MHz, $CDCl_3$): δ 855 (d, *J* = 1.1 Hz, 1H), 7.90 (s, 1H), 7.57 (d, *J* = 8.8 Hz, 2H), 7.09 (d, *J* = 1.5 Hz, 1H), 6.99 (d, *J* = 8.8 Hz, 2H), 4.66 (dt, *J* = 47.0, 5.7 Hz 2H), 4.17 (t, *J* = 6.1 Hz, 2H), 3.76 (s, 3H), 3.74–3.70 (m, 2H), 3.69–3.53 (m, 8H), 3.50–3.44 (m, 4H), 3.35–3.32 (m, 1H), 3.11–3.06 (m, 2H), 2.29–2.12 (m, 2H), 1.57–1.29 (m, 15H). $^{13}C$ NMR (150 MHz, $CDCl_3$, 60℃): *δ* 175.14, 170.29, 163.51, 161.22, 161.14, 158.59, 157.77, 156.1, 148.20, 130.05, 126.89, 115.98, 114.67, 100.01, 80.51, 79.18, 70.56, 70.39, 70.14, 69.72, 64.13, 55.53, 41.20, 40.57, 39.05, 38.65, 34,96, 30.51, 30.04, 28.55, 23.01.

Compound **7**: To a solution of compound **22** (3.0 mg, 4.0 μmol, 5 eq.) in dry DMSO (120 μL), Alexa Fluor™ 647 NHS ester (1.0 mg, 0.8 μmol, 1.0 eq.) and DIPEA (1 μL, 5.7 μmol, 7.2 eq.) were added. After stirring at RT overnight, the solvent was removed by lyophilization. The residue was dissolved in TFA/$CH_2Cl_2$ (1/1) (1 mL) and stirred at RT for 30 min. The solvent and TFA was removed by evaporation and the residue was purified by reversed-phase HPLC with a linear gradient of 15-30 % $CH_3CN$-0.1% TFA (20 min) to afford **7** as a blue film. HR-ESI-MS: calcd. for $C_{66}H_{87}FN_{10}O_{18}S_5$ $[M-3H+2Na]^-$ = 1529.4353, obsd. = 1529.4491. calcd. for $[M-3H+Na]^{2-}$ = 753.2231, obsd. =753.2218.

**Preparation of Nb381-Ax647 (Probe 4):**

**Preparation of Plasmid Construct**

For chemical dye conjugation, tandem GS linker and the Cys residue were introduced in front of the His tag sequence using primers, 5'- GGG TCC TGT CAC CAC CAT CAC CAT CAC GAA CCT GAA GCC -3'/ 5'- GCC ACC GCC GCT GCC GCC ACC CCC GGA -3' and Q5 Site-Directed Mutagenesis Kit (New England Biolabs) to confer pMESy4-PelB-Nb381-$(GS)_2$-Cys-6His (Supplementary Figure 13). The DNA sequence of the plasmid construct was verified by dye-terminator DNA sequencing (Genewiz).

**Protein Expression and Purification**

The plasmid pMESy4-PelB-Nb381-$(GS)_2$-Cys-6His was transformed into a BL21(DE3) strain using the standard heat-shock protocol. The transformed BL21(DE3) cells was grown in a Luria-Bertani (LB) medium containing 50 μg/mL kanamycin for 16 h at 37 ºC as a starting culture. The starting culture (1 mL) was transferred into a fresh Terrific broth (TB) medium and incubated at 37 ºC until an OD at 600 nm of ~ 0.8 was reached at which time, cells were induced with 1 mM IPTG. Cells were incubated for another 5 h at 37 °C, pelleted by centrifugation, and frozen at -20 °C overnight. Cells were resuspended in a PBS buffer (Thermo Fisher) on ice bath and lysed using standard sonication protocols. The soluble fractions were loaded onto TALON beads (Takara Bio) pre-equilibrated with PBS. After washing the column with 10 volumes of PBS, the target protein was eluted with PBS containing 500 mM imidazole. The excess imidazole was removed by dialysis using 3.5 K membrane filter (Spectrum Laboratories Inc.). The purified protein was concentrated using 10 K Amicon centrifuge filter (Millipore). Prior to protein-dye conjugation, purified Nb381-$(GS)_2$-Cys recombinant protein was treated using endotoxin removal spin column (Thermo Fisher). The purity and MW of protein were confirmed by SDS-PAGE analysis.

**Alexa Fluor 647 maleimide conjugation**

To a solution of Nb381-$(GS)_2$-Cys in PBS (300 μM, 500 μL) containing 10 mM TCEP, Alexa Fluor 647 maleimide (100 μM) was added and the reaction mixture was incubated at 24 °C overnight under the dark. The nanobody-dye conjugate was purified using a TOYOPEARL HW-40F size-exclusion column (TOSOH) and Hi-Trap QHP anion exchange column (GE Healthcare). The purity and MW of nanobody-dye conjugate was confirmed by SDS-PAGE, in Gel fluorescence and MALDI-TOF MS analysis (Supplementary Figure 14 and 15).

**Expression of receptors in HEK293T cells.**

HEK293T cells (ATCC) were cultured in Dulbecco's modified Eagle's medium (DMEM) supplemented with 10% fetal bovine serum (Sigma Aldrich), penicillin (100 units $mL^{-1}$), streptomycin (100 μg $mL^{-1}$) and amphotericin B (250 ng $mL^{-1}$) and incubated in a 5% $CO_2$ humidified chamber at 37 °C. For expression of each receptor, HEK293T cells ($2.0 \times 10^5$ cells) plated on a 3.5-cm dish (Corning) were transfected with each expression vector for HA-tag fused mGlu1[S6] using Lipofectamine 2000 (Invitrogen) according to the manufacturer's instructions.

**Confocal imaging of HEK293T cells stained with an imaging probe.**

After 24 h of transfection, the cells were dissociated by treating with TrypLE Express (Gibco) and re-seeded on 35 mm glass-bottom dishes (Iwaki) pretreated with poly-L-lysine. After 24 h of the re-seeding, the cells were washed twice with PBS(–) (Wako) then 100 μL of PBS(-) containing a probe (2 μM) was added into the cell cultured dish. After incubation at RT for 5 min, 100 μL of 4% PFA/PBS(–) was added. After incubation at RT for 30 min, cells were washed with DMEM at 37℃. After HA-tag staining (DyLight550 anti-HA tag, ab117502), fluorescence imaging of the cells was performed using CLSM equipped with a 63×, NA = 1.40 oil objective, and GaAsP detector. Fluorescence images were acquired using the 561 nm excitation for DyLight550 and the 633 nm excitation for Alexa Flour 647 derived from a white laser.

**Determination of an apparent affinity constant of the probes.**

The HEK293T cells transiently transfected with HA-tag fused mGlu1 on a 35 mm glass-bottom dishes were washed twice with PBS(–), and then 150 μL of PBS(–) solutions containing a probe at the different concentrations were added to the dishes. After incubation at RT for 5 min, 150 μL of 4% PFA/PBS(–) was added to each dish. After incubation at RT for 30 min, cells were washed twice with PBS(–). After HA-tag staining, the fluorescence imaging was performed. The fluorescence intensity of a probe bound to mGlu1 and HA-staining signal at each concentration was determined by enclosing the regions containing a cell membrane with ROIs and $F = F_{probe}/F_{HA}$ was calculated for each cell ($F_{probe}$ is a fluorescence intensity of probe and $F_{HA}$ is a fluorescence intensity of HA-staining signal) (n = 8). After plotting the averaged and background-subtracted value against probe concentrations, the apparent dissociation constant ($K_d$) was determined by fitting with the theoretical logistic equation.

$$F = F_{sat} / (1 + (\log [\text{probe}] / \log K_d)^n)$$

$$\log n = h$$

$F_{sat}$ is a saturation point.

$F_0$ is a fluorescence intensity in the absence of probe.

$F_{background}$ is an averaged fluorescence intensity of background.

h is a Hill's coefficient

**Perfusion fixation and brain slices preparation.**

Experiments were conducted according to the literature.[S7] Briefly, under the deep anesthesia with isoflurane, mice were perfused transcardially with ice-colded 4% formaldehyde/PBS(–) (pH 7.4) (60 mL). The mouse brain samples were fixed with 4% PFA at 4 °C overnight. After washing with PBS(–) (x3), the brain samples were immersed into 30% sucrose/PBS. The brain slices were prepared using a cryostat (Leica, CM-1950).

For the immunostaining, the brain slices were permeabilized with PBS(–) containing 0.1% triton X-100 for 15 min and blocked with 10% normal goat serum (NGS) in PBS(–) containing 0.1% triton X-100 for 30 min. Then, primary antibody reaction was conducted with rabbit anti-mGlu1 (Frontier Institute, MSFR104030), rabbit anti-Calbindin (Frontier Institute, CB-28kD), rabbit anti-DRD2 (Frontier Institute, AB_2571596), rabbit anti-5HT2A (ImmunoStar, 24288), rabbit anti-GluD2 (SIGMA, HPA056253), guinea pig anti-Shank2 (Synaptic systems, 162 204) or mouse anti-Synaptophysin (abcam, ab8049) in PBS(–) containing 0.1% triton X-100 at 4 °C overnight. For the immunostaining of GluA2, the brain slice (15 μm thickness) was attached to glass slide and activated with antigen retrieval reagent ImmunoSaver (FUJIFILM Wako) at 80ºC for 20 min. After blocking with 10% NGS in PBS(–) containing 0.1% triton X-100 for 30 min, primary antibody reaction was conducted with mouse anti-GluA2 (Merck, MAB397). Secondary antibody reaction was conducted with a goat anti-guinea pig IgG H&L (Alexa Fluor® 405) (ab175678, for Shank2), goat anti-rmouse IgG H&L (Alexa Fluor® 488) (ab150113, for Synaptophysin), goat anti-rabbit IgG H&L (Alexa Fluor® 488) (ab150077, for mGlu1, calbindin, DRD2, 5HT2A, and GluD2) or goat anti-mouse IgG H&L (Alexa Fluor® 594) (abcam, ab150116, for GluA2) in PBS(–) containing 0.1% triton X-100 at r.t. for 1 h.

**Clearing of mouse brain tissue with 3DISCO protocol.**

Experiments were conducted according to the literature.[S8] The fixed brain sample with 4%

PFA/PBS(–) was treated in the mixture of THF/$H_2O$ (vol./vol.) at a series of concentrations (50, 70, 80 (each for 12 h) and 100% (12 h × 3) with shaking at RT on a turning table under the dark. Finally, the brain sample was immersed in dibenzyl ether for refractive index matching for 1–2 day before CLSM imaging.
